# The stationary phase-specific sRNA *fimR2* is a multifunctional regulator of bacterial motility, biofilm formation and virulence

**DOI:** 10.1101/2022.02.17.480891

**Authors:** Nicole Raad, Disha Tandon, Siegfried Hapfelmeier, Norbert Polacek

**Affiliations:** Department of Chemistry, Biochemistry, and Pharmaceutical Sciences, University of Bern, Bern, Switzerland; Graduate School for Cellular and Biomedical Sciences, Bern, Switzerland; Institute for Infectious Diseases, University of Bern, Bern, Switzerland

**Keywords:** sRNA, ncRNA, biofilm formation, type 1 pilus, type III secretion system, virulence, motility, stationary phase, RNase E, CsrA

## Abstract

Bacterial pathogens employ a plethora of virulence factors for host invasion, and their use is tightly regulated to maximize infection efficiency and manage resources in a nutrient-limited environment. Here we show that during *Escherichia coli* stationary phase the small non-coding RNA *fimR2* regulates fimbrial and flagellar biosynthesis at the post-transcriptional level, leading to biofilm formation as the dominant mode of survival under conditions of nutrient depletion. *fimR2* interacts with the translational regulator CsrA, antagonizing its functions and firmly tightening control over motility and biofilm formation. Generated through RNase E cleavage, *fimR2* regulates stationary phase biology independently of the chaperones Hfq and ProQ. The *Salmonella enterica* version of *fimR2* induces effector protein secretion by the type III secretion system and stimulates infection, thus linking the sRNA to virulence. This work reveals the importance of bacterial sRNAs in modulating various aspects of bacterial physiology including stationary phase and virulence.

**Highlights:**

- *fimR2* expression causes biofilm formation and alters bacterial outer membrane architecture
- *fimR2* modulates CsrA activity and sequesters it from its targets
- The *Salmonella fimR2* variant is functional in *E. coli*
- *fimR2* is generated through RNase E processing and enhances infectivity

## Introduction

The ability of bacterial pathogens to infect and invade a host critically relies on the presence of adequate conditions promoting the survival of the pathogen within the said host, and the available arsenal of virulence factors allowing the execution of the various infectious steps. Once initial contact has been established and infection initiated, pathogens have to navigate their way through the various assaults of the host immune response, fighting back at times and defending their grounds at others, to reach regions with more resources. In this process, attacking bacterial pathogens express and employ a variety of virulence factors that either harm the host, such as toxins, or promote tissue invasion, such as adhesins (Wilson et al., 2002). For instance, several Gram-negative pathogens such as enterohaemorrhagic *Escherichia coli* (*EHEC*), enteropathogenic *E. coli* (*EPEC*), enteroinvasive *E. coli* (*EIEC*), and *Salmonella enterica* serovar Typhimurium employ the type III secretion system (T3SS) to deliver a wide range of effector proteins directly into the cytoplasm of their target cells (Kaper et al., 2004, Galán, 2021) and these effectors cause cytotoxicity and manipulate the actin cytoskeleton to promote bacterial invasion. Uropathogenic *E. coli* (*UPEC*) use their type 1 pili (T1P) to adhere to and invade bladder epithelial cells (Martinez et al., 2000) while *S. enterica* uses the same pili to bind microfold cells within the intestinal lumen (Hase et al., 2009).

Which virulence factors to use and when to employ them are two important aspects for microbial infections not only to ensure a successful attack, but also to minimize the loss of resources for the pathogen in an unpredictable and harsh environment (Sy and Tree, 2021). For these reasons, bacterial virulence factors are under tight regulation via multiple mechanisms acting in concordance and have been refined across long stretches of evolutionary history. CsrA is a translational regulator that governs the expression of T3SS effectors in both *S. enterica* and *E. coli.* For instance, CsrA downregulates *hilD*, the SPI-1 (Salmonella pathogenicity island 1) encoded regulator, and by doing so, inhibits the expression of multiple HilD-dependent T3SS effectors (Martínez et al., 2011). In *EPEC*, CsrA induces flagellar motility and the expression of some T3SS effectors involved in Attaching/Effacing (A/E lesions) (Bhatt et al., 2009). Both *Salmonella* and *E. coli* regulate the expression of T1P through phase variation (Henderson et al., 1999), allowing some members of isogenic populations to display the pilus and others to retract it. However, the precise modes of regulations are far more intricate. In *Salmonella,* the expression of T1P is controlled by regulatory proteins FimZ, FimY, and FimW that affect expression from the single promoter of the *fimAICDHF* operon encoding T1P (Kolenda et al., 2019). In *E. coli,* the promoter of *fimAICDFGH-*encoding T1P lies within an invertible DNA region (Abraham et al., 1985) called *fimS*, and recombinases FimB and FimE govern its orientation: FimB mediates the inversion in both ON (expression) and OFF (repression) orientations while FimE is biased to the OFF inversion (McClain et al., 1991). This complex mode of regulation in both pathogens is further controlled by additional transcriptional regulators [Reviewed in Schwan (2011) and in Kolenda et al. (2019), tightening the rope around the expression of this costly virulence factor (Sterzenbach et al., 2013).

Biofilm formation is another mode employed by bacterial pathogens to assist in host infection. This multicellular lifestyle benefits bacteria in multiple ways, the most important of which is resistance to external assaults or internal stresses. Indeed, biofilms have been linked to the establishment of recurrent infections and bacterial persistence (Naziri et al., 2021). UPEC can form resilient biofilms through the coordinated expression of several outer membrane appendages, including pili, fimbriae, flagella, and curli (Kaper et al., 2004, Pratt and Kolter, 1998). The T1P plays important roles in this context, allowing *UPEC* to form biofilms on both abiotic surfaces such as medical supplies and urinary catheters (Wang et al., 2018), and in host tissues, such as the bladder epithelium where specialized biofilms called intracellular bacterial communities (IBCs) cause chronic infections (Wright et al., 2007).

Recently, *trans*-encoded small non-coding RNAs (sRNAs) have emerged as potent regulators of gene expression in various bacterial species (Hör et al., 2020). By partially base pairing with various target mRNAs and regulating their expression at the post-transcriptional level, these non-coding RNAs (ncRNAs) govern various aspects of bacterial physiology (Nitzan et al., 2017), with various roles described in bacterial virulence and biofilm formation (Sy and Tree, 2021, Chambers and Sauer, 2013). For instance, *EHEC GlmY* and *GlmZ* sRNAs promote the formation of A/E lesions through induction of the T3SS effector EspFu, and diminish the expression of other T3SS effectors (Gruber et al., 2014). In *Salmonella*, the sRNA *PinT* coordinates the expression of *SPI-1* and *SPI-2* (Salmonella pathogenicity islands 1 and 2) T3SS virulence factors (Westermann et al., 2016) allowing the pathogen to switch from its invasive to its persistent infective mode. Salmonella *CpxQ* sRNA inhibits *fimA* expression through direct base-pairing with the *fimA* mRNA, controlling T1P expression under conditions of membrane stress (Chao and Vogel, 2016). In *E. coli*, various sRNAs have been shown to affect T1P expression through the direct inhibition of *fimA* or *fimB*, such as the sRNAs *RybB* and *MicA* (Bak et al., 2015). These sRNAs often require the RNA chaperone Hfq for generation, stability, and function (Santiago-Frangos and Woodson, 2018). In recent years, similar roles have been ascribed to ProQ, a FinO-family member (Smirnov et al., 2017). While these chaperones are critical for sRNA stability and function, examples of sRNA independence exist in Gram-positive species. For example, *Mycobacterium tuberculosis 6S* sRNA acts independently of chaperone proteins to coordinate cell division and DNA replication (Mai et al., 2019), teasing the possibility that chaperones can be dispensable for some regulatory circuits.

Previously, our deep sequencing analysis of growth phase-dependent sRNAs in *E. coli* revealed the abundant expression of an sRNA, *sRNA_35*, in the stationary phase (Raad et al., 2021). This sRNA has also been recognized in previous sequencing studies (Kawano et al., 2005, Ghosal et al., 2015). Here, we report the first functional characterization of this 35-nucleotide sRNA that we rename to *fimR2* for *fim* operon-derived sRNA as it derives from the *fimA-fimI* intergenic region in *E. coli* and *S. enterica.* This operon has been shown by others to encode another sRNA, dubbed *fimR*, which however derives from a different location within the *fim* operon and is thus completely independent of *fimR2* (Pichon et al., 2012). *fimR2* overexpression in exponential phase, a growth phase during which this sRNA is normally undetectable, significantly causes biofilm formation, alters the outer membrane architecture, and switches the bacterial population to stationary phase. Through various genetic and biochemical approaches, we show that these phenotypes are due to the *fimR2-*mediated regulation of various targets involved in stationary phase biology and motility, directly through base-pairing with target mRNAs, and indirectly, through the sequestration of the translational regulator CsrA. Interestingly, overexpression of the *Salmonella* variant, *fimR2S*, in *E. coli* causes biofilm formation, potentiates the invasiveness of *S. enterica,* and promotes the expression of a T3SS-chaperone. In sharp contrast to most bacterial *trans*-acting sRNAs, we show that *fimR2* acts independently of RNA matchmakers such as Hfq and ProQ and accumulates through the RNase E-dependent cleavage of *fimAICDFGH*. We thus propose an example of a *trans*-acting sRNA in Gram-negative bacteria that does not require an RNA chaperon for its function. This work positions *fimR2* as a master regulator of gene expression under stationary phase, employing multitasking to coordinate biofilm formation and virulence.

## Results

### *fimR2* is abundantly expressed in various *E. coli* strains in a phase-dependent manner

We previously sequenced the small *E. coli* transcriptome and identified several sRNAs that are differentially expressed during stationary phase (Raad et al., 2021). One of these sRNA candidates is a 35-nucleotide-long ncRNA (initially dubbed *sRNA_35*) whose sequence maps to the intergenic space between *fimA* and *fimI*. These two protein-coding genes are members of the *fim* operon encoding the components of T1P. Due to its genomic context, we renamed *sRNA_35* to *fimR2* for *fim operon*-derived sRNA. We were intrigued by the abundant expression of *fimR2* in stationary phase (**Figure S1A**) knowing that *fimAICDFGH* is not expressed in this phase as previously reported (Dove et al., 1997). Indeed, we confirmed that the phase-dependent expression of *fimR2* is coupled to the concomitant downregulation of *fimAICDFGH* expression (**Figure S1A**).

The conservation of *fimR2* across various *E. coli* strains and within enteric bacteria (**Figure S1B**) suggests a role for this sRNA resembling that seen in the domesticated laboratory strain, K12. Thus, we evaluated the expression of this sRNA by northern blotting in five extended spectrum beta-lactamases (ESBL)-producing *E. coli* strains derived from healthy donors (Moor et al., 2021) (**Figure S1C**). All five strains retain the genomic sequence of *fimR2*, while four strains expressed the sRNA in stationary phase to varying extents in comparison to the K12 laboratory strain. Extending our experiments also to pathogenic *E. coli* strains revealed an abundant expression of *fimR2* in a verrotoxin-producing *E. coli* (VTEC) and two UPEC strains in a phase-dependent manner, peaking in late stationary phase (**Figure S1D**). Taken together, these findings suggest that the *fimR2* abundant expression in stationary phase is a global occurrence that likely plays important roles in this context.

### *fimR2* is an RNase E-dependent sRNA

The abundant expression of *fimR2* in stationary phase of *E. coli* raises questions about the mechanism of its biogenesis. As *fimR2* is expressed in stationary phase, its expression could be governed by the alternative sigma factor RpoS which regulates the transcription of several genes in stationary phase (Hengge-Aronis, 2002). However, northern blot analysis showed an increase in abundance of *fimR2* following *rpoS* deletion (**Figure S2A**) suggesting that this sRNA is not a primary transcript. Furthermore, expression of *fimR2* was not diminished by a *fimA* deletion (**Figure S2B**) which abolishes a predicted internal transcription start site (TSS) (Thomason et al., 2015), suggesting that *fimR2* expression does not depend on this TSS for expression.

These findings indicated *fimR2* as a processing product from the *fimAICDFGH* primary transcript. If this is the case, the expression of *fimAICDFGH,* which is governed by the two recombinases FimB and FimE (McClain et al., 1991), is a prerequisite for *fimR2* biogenesis. Knockout of FimB, which allows both ON and OFF inversions to take place, abolished *fimR2* expression in stationary phase (**Figure S2C**). In the absence of FimB, FimE directs *fimS* to the OFF orientation and therefore *fimAICDFGH* cannot be expressed and *fimR2* not processed. However, the knockout of the phase-OFF-specific recombinase FimE leads to a prominent upregulation of *fimR2* in both exponential and stationary phase (**Figure S2C**). Additionally, *fimR2* processing intermediates with various sizes accumulated exclusively in exponential phase upon *fimE* deletion, but not in stationary phase (**Figure S2C**). In the absence of FimE, FimB inverts *fimS* in both orientations without being antagonized by FimE, causing prominent expression of *fimAICDFGH* (data not shown). These experiments suggested that the biogenesis of *fimR2* necessitates the prior expression of *fimAICDFGH* and that *fimR2* is most likely cleaved from this transcript by an endonuclease specifically in stationary phase.

We next sought to determine the identity of the RNase(s) responsible for *fimR2* processing and release in stationary phase. For this purpose, RNA was extracted from several RNase deletion strains and *fimR2* expression was evaluated by northern blotting. Knockout of RNase I, PNPase, RNase G, RNase R, RNase HI or RNase HII did not affect *fimR2* levels during stationary phase (**Figure S2D**). Similarly, a mutation that impairs RNase III function (Dasgupta et al., 1998) did not affect *fimR2* abundance (**Figure S2E**). These experiments suggested that these nucleases are not involved in *fimR2* biogenesis.

A key endoribonuclease that is involved in mRNA turnover in stationary phase and in sRNA biogenesis is the endoribonuclease E (RNase E) (Hör et al., 2020). RNase E cleavage generates RNA molecules with a monophosphate at the 5’-end (Chao and Vogel, 2016). To determine the nature of *fimR2* 5’-end, we subjected total RNA to a terminator exonuclease (TEX) treatment that preferentially degrades RNA molecules with a 5’-monophosphate, such as rRNA. *fimR2* was sensitive to TEX treatment, but not an *in vitro* transcribed tRNA fragment (tRF) with a 5’-triphosphate (**Figure 1A**), suggesting that *fimR2* is processed and retains a 5’-end signature typical for RNase E cleavage. To investigate the putative involvement of RNase E in *fimR2* processing, an RNase E-temperature sensitive strain was employed that expresses a fully active RNase at 30°C and an RNase with impaired function at 43°C (Babitzke and Kushner, 1991). Upon switching the stationary phase-bacterial culture from the permissive to the non-permissive temperatures, *fimR2* was depleted in *rne3071^-ts^* strain but not in the K12 strain (**Figure 2B**). Simultaneously, the precursor of the RNase E-dependent sRNA *ArcZ* (Updegrove et al., 2018), accumulated in the *rne3071^-ts^* strain at the non-permissive temperature. This confirmed the role of RNase E in *fimR2* processing. To further corroborate these findings, we used the purified catalytic domain of RNase E (Callaghan et al., 2005) in an *in vitro* cleavage assay. For this, a *fimR2* precursor containing extensions on both ends was *in vitro* transcribed and then RppH-treated to generate RNase E cleavage templates containing a monophosphate at the 5’-end (Jiang and Belasco, 2004). Incubation of the precursor with increasing amounts of RNase E led to the cleavage of the precursor to an intermediate size of 40 nt (**Figure 2C**).

**Figure 1:**
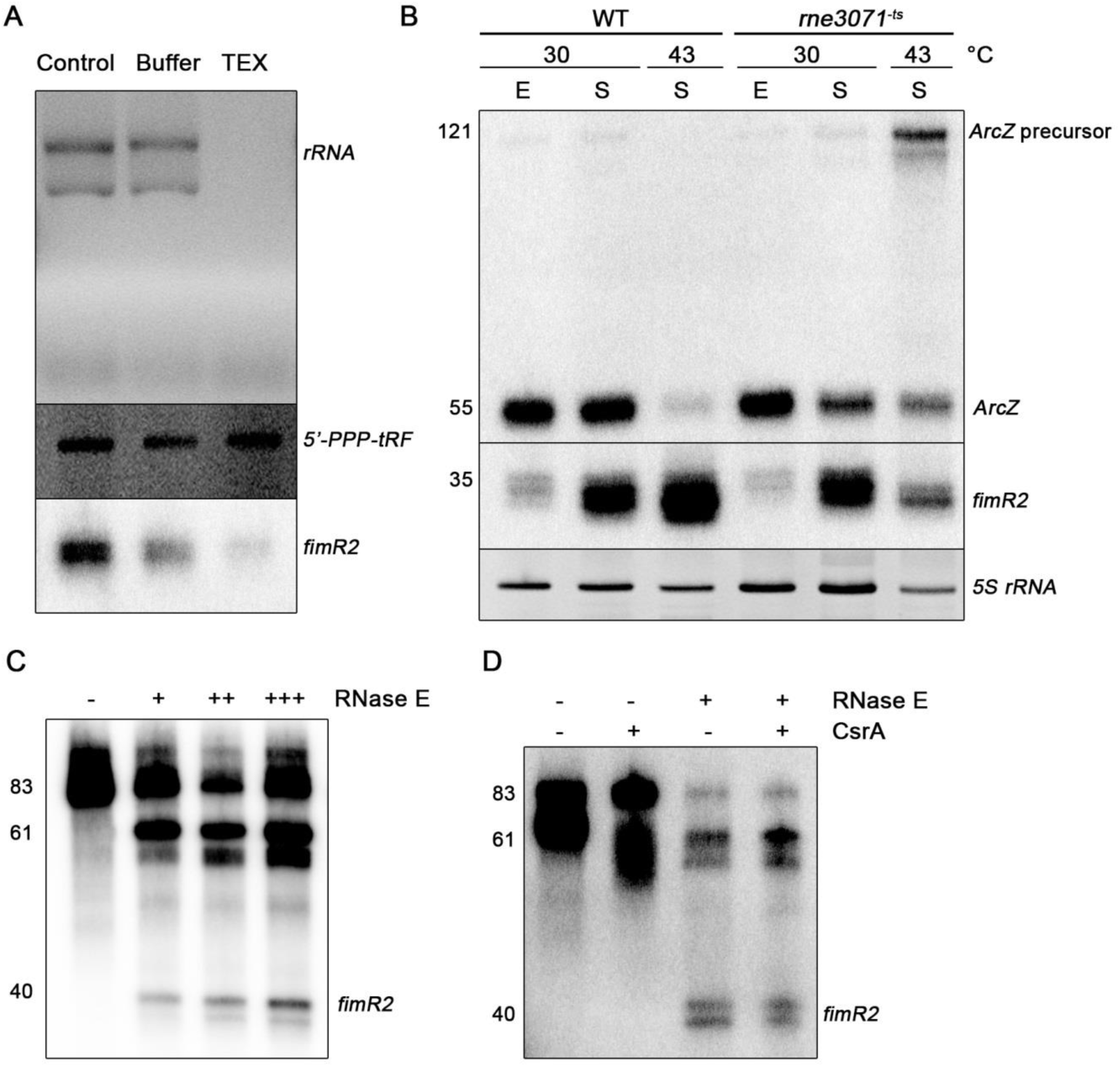
*fimR2* is processed by RNase E. **A.** Ethidium bromide staining of ribosomal RNA and 5’-triphosphorylated tRNA fragment, and northern blot analysis of *fimR2* upon TEX treatment. Total RNA samples were untreated (control), incubated with buffer, or with TEX. **B.** Northern blot analysis of *fimR2* and *ArcZ* sRNAs expression in WT (wild-type) and *rne3071^-ts^* (RNase E temperature-sensitive) strains. E and S denote total RNA samples extracted from exponential phase and stationary phase, respectively, and from incubations at the indicated temperatures. Ethidium bromide staining of *5S rRNA* is shown as a loading control. **C.** Northern blot analysis of *fimR2* following *in vitro* cleavage of a 5’- and 3’-extended precursor with (+) and without (–) increasing amounts of RNase E. **D.** Northern blot analysis of *fimR2* following *in vitro* cleavage of a 5’- and 3’-extended precursor with (+) or without (–) RNase E and CsrA. In B-D the size of the different RNA molecules is indicated on the left.

**Figure 2:**
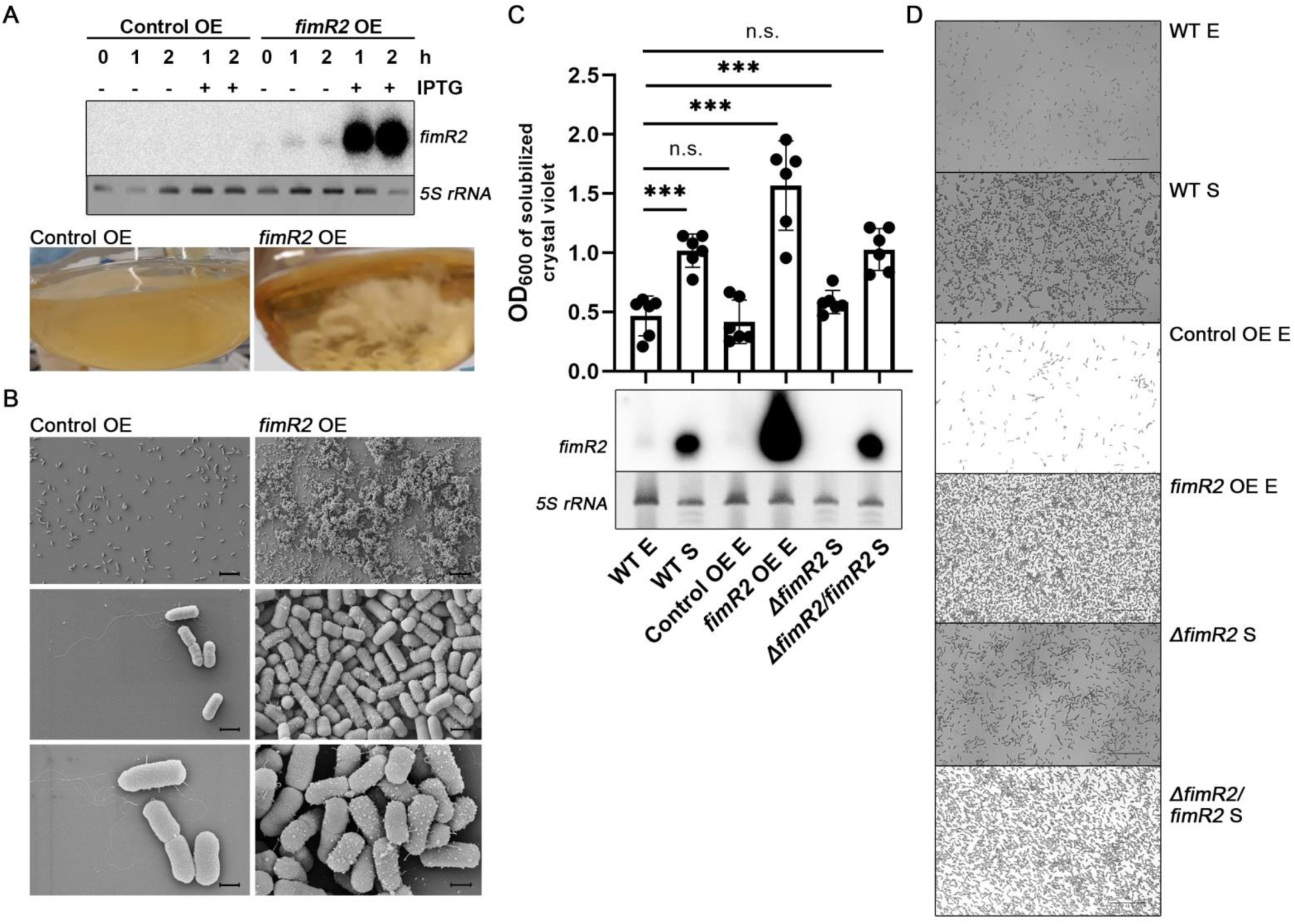
*fimR2* regulates biofilm formation. **A.** Northern blot (top) of *fimR2* expression in exponential phase following overexpression (OE) of the sRNA via IPTG induction. Pictures of resulting cultures are shown on the bottom. Control OE refers to the expression of a random short RNA sequence. Ethidium bromide staining of 5S *rRNA* is shown as a loading control. **B.** Scanning electron micrographs of coverslip-formed biofilms of Control and *fimR2* OE strains. Scale bars = 10 μm (top), 1 μm (middle), and 500 nm (bottom). **C.** Northern blot of *fimR2* expression in WT (wild-type), Control OE, *fimR2* OE (*fimR2* overexpression), *ΔfimR2* (*fimR2* deletion), and *ΔfimR2*/*fimR2* (*fimR2* complementation) strains. Samples are shown from exponential (E) and stationary (S) phase. Ethidium bromide staining of 5S rRNA is shown as a loading control. **D.** Quantitative biofilm assay showing mean + SD OD_600_ of solubilized crystal violet-staining from six biological replicates of strains and conditions in C. Unpaired two-tailed t-test with Welch’s correction was used to determine significance with n.s and *** showing not significant and significant results, respectively. The p-values are from bottom to top 0.6369, 0.0003, 0.0001, 0.0002, and 0.9202, respectively. **E.** Micrographs of air-liquid phase biofilms stained with crystal violet, under conditions mentioned in C. Scale bar = 25 μm.

Taken together, these findings suggest that RNase E governs *fimR2* biogenesis from the parental *fimAICDFGH* transcript in stationary phase. The appearance of the 40-nt intermediate in the *in vitro* cleavage assay suggested *fimR2* to be trimmed to its final size by another yet to be identified RNase, or that *in vivo*-derived *fimR2* is post-transcriptionally modified.

### *fimR2* promotes stationary phase-dependent biofilm formation

What kind of functions is the RNase E-processed *fimR2* involved in? We previously reported that the ectopic overexpression of this sRNA in exponential phase, its alienated phase, upregulates the expression of *rpoS* (Raad et al., 2021), the alternative RNA Polymerase sigma factor governing gene expression in stationary phase. This suggested that the sRNA is important for stationary phase-specific phenomena. To gain insight into possible *fimR2* functions, we ectopically expressed this sRNA in an inducible manner during exponential phase (Raad et al., 2021). Strikingly, *fimR2* expressing strains started to dramatically aggregate in culture following 1 h of overexpression at static conditions, and beyond 2 h at shaking conditions, coupled to the clearing of the LB medium, a phenotype with no counterpart in the control strain (**Figure 2A**). Furthermore, the aggregated cellular suspension forms resistant biofilms at the bottom and edges of the flask, suggesting that *fimR2* overexpression causes biofilm formation. To inspect this phenotype further, we performed scanning electron microscopy (SEM) of the *fimR2* overexpressing strain grown in microtiter plates mounted with coverslips. SEM revealed significant crowding of the coverslip with multiple embedded cells for the *fimR2* overexpression strain, and only a modest bacterial population for the control strain, which overexpressed an unrelated RNA sequence (**Figure 2B**). We further corroborated these findings using quantitative biofilm assays and a gentler qualitative biofilm assay of bacterial cells grown on coverslips. Both approaches confirm that *fimR2* overexpression causes biofilm formation (**Figure 2C, D**). As bacterial cells in stationary phase often form biofilms (Markova et al., 2018), we next inspected the biofilm forming potential of wild-type (WT) and *fimR2* genomic deletion *(ΔfimR2)* strains in stationary phase. Indeed, both WT and *ΔfimR2* strains formed biofilms in stationary phase (**Figure 2C, D**) while *ΔfimR2* showed markedly reduced biofilm formation potential compared to the WT strain. Genetic complementation of *fimR2* deletion (*ΔfimR2/fimR2*) through ectopic expression of this sRNA restores biofilm formation to WT levels (**Figure 2C, D**). Taken together, these experiments suggested that *fimR2* is an important contributor to biofilm formation in stationary phase.

To further validate the involvement of *fimR2* in this phenotype, we carried out mutagenesis studies. A careful observation of the aligned *fimR2*-like sequences in the *fimA-fimI* region from various *E. coli* strains and other enteric bacteria suggests that the sequence is highly conserved, with nucleotides 18 to 31 having the least variations (**Figure S1B**). MFold analysis predicted this region to forming a stem-loop secondary structure (**Figure S3A**). Mutations that individually disrupt the stem or the loop of this hairpin destabilized the sRNA (**Figure S3B**) and consequently eliminated the biofilm formation (**Figure S3C, D**). A *fimR2* compensatory mutant that allows the re-establishment of the stem-loop structure was stably expressed (**Figure S3B**) and functional in eliciting biofilm formation albeit to a lesser extent than the wild-type sequence (**Figure S3C, D**). Interestingly, the *fimE* deletion strain, a strain constantly transcribing the *fimAICDFGH operon,* phenocopied *fimR2* overexpression in exponential phase and caused prominent biofilm formation (**Figure S2F, G).** We conclude that this secondary structure likely exists *in vivo* and contributes to *fimR2* stability, and thus its ability to propagate biofilm formation.

### *fimR2* alters bacterial outer membrane architecture

The dramatic biofilm formation phenotype seen upon *fimR2* expression is expected to be accompanied with an altered gene expression pattern that will remodel the outer membrane architecture and impact proteinaceous appendages displayed to the outer milieu, such as flagella, fimbriae, and pili. To evaluate this hypothesis, we used scanning electron micrography of *E. coli* bacterial pellets from planktonic growth stages. SEM revealed that bacterial cells derived from exponential phase of growth were elongated and displayed flagella, fimbriae, and pili (**Figure 3A**). In contrast, bacterial cells in stationary phase were more ovoid and naked, displaying curli fimbriae and PGA (poly-β-1,6-N-acetyl-d-glucosamine) (Boehm et al., 2009) (**Figure 3B**). Strikingly, *fimR2* overexpressing strains in exponential phase displayed pronounced aggregation and PGA synthesis and adopted a stationary phase-like morphology while the control overexpressing strains resembled exponentially growing cells (**Figure 3C, D**). *ΔfimR2* and *ΔfimR2/fimR2* strains significantly diverged in their morphologies from one another, although both strains were in stationary phase. *ΔfimR2* exhibited exponential phase-like morphologies with extensive flagellation, while *ΔfimR2/fimR2* mimics wild-type stationary phase physiology (**Figure 3E, F**). The characteristic morphologies of cells in exponential and stationary phase are in agreement with what has been previously reported (Serra et al., 2013), showing that our distinct results were not artifacts of cell fixation. These SEMs thus showed that *fimR2* expression and *fimR2* deletion resulted in opposing morphologies on the extreme ends of a spectrum that separates stationary from exponentially growing cells, with control overexpressing and *fimR2* compensation strains lying in the middle and behaving like their wild-type counterparts in their respective stages. In this manner, *fimR2* appeared to trigger a significant outer membrane architecture remodeling, a critical process in the transition to stationary phase.

**Figure 3:**
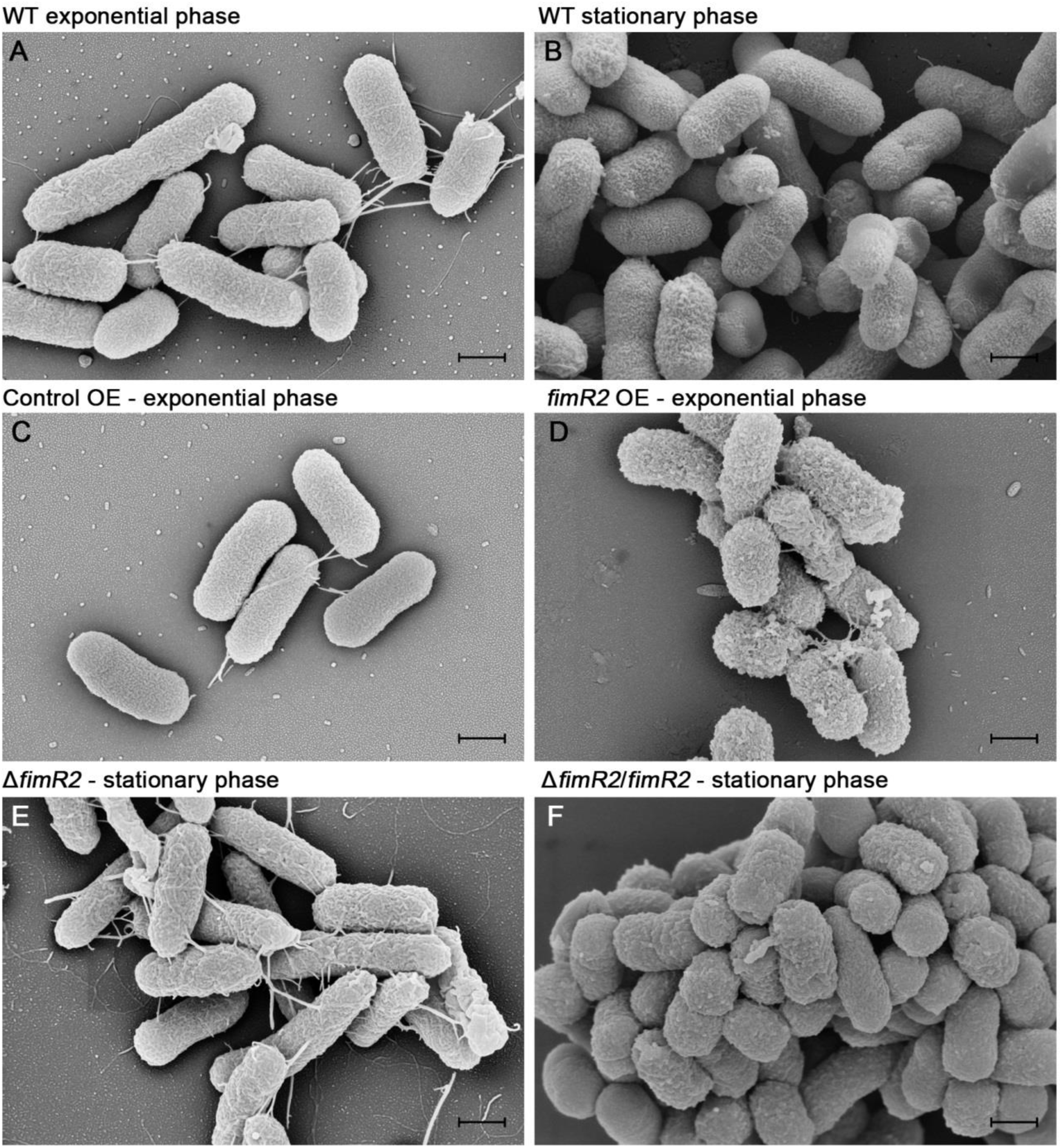
*fimR2* alters bacterial outer membrane architecture. Scanning electron micrographs of *E. coli* K12 strains in **A.** WT exponential phase, **B.** WT stationary phase, **C.** Control OE (Control overexpression) in exponential phase, **D.** *fimR2* OE (*fimR2* overexpression) in exponential phase, **E.** *ΔfimR2* in stationary phase, and **F.** *ΔfimR2*/*fimR2* in stationary phase. Scale bar = 500 nm.

### *fimR2* regulates cell motility and stationary phase biology

To test the hypothesis that *fimR2* binds putative mRNA targets via its predicted single-stranded region (**Figure S3A**), a bioinformatic target prediction tool was employed (Wright et al., 2014). Guided by the observed *fimR2*-dependent phenotype (**Figures 2 and 3**), a subset of the predicted 200 targets in K12 strain were selected for experimental validation. Multiple predicted targets (*fliJ, fliG*, *fliI,* and *fliR*) are mRNAs coding for flagellar biosynthesis proteins (**Figures 4A** **and S4A-C**) (Minamino and Namba, 2004) three of which (*fliJ*, *fliG*, and *fliI*) are part of the same operon (Fitzgerald et al., 2014). Additionally, one of the top candidates, *hofQ*, (**Figure S4D**) is a poorly characterized importer of exogenous DNA to be used in catabolic reactions that provide more nutrients to the starved cells in stationary phase (Palchevskiy and Finkel, 2006). First, we evaluated the expression pattern of these putative targets by RT-qPCR analysis under different conditions. The flagellar biosynthesis operon and *fliR* were downregulated upon *fimR2* overexpression during exponential growth, a pattern resembling the canonical expression of these messages in stationary phase of growth (**Figures 4B** **and S4A-C**). *hofQ* was upregulated upon *fimR2* overexpression during exponential phase (**Figure S4D**). The genomic deletion of *fimR2* (*ΔfimR2)* perturbed the stationary phase expression pattern of all candidates while in the *fimR2* complementation (*ΔfimR2/fimR2)* strain the stationary phase-specific expression was restored (**Figures 4B** **and S4A-D).** These findings suggested these mRNAs as genuine *fimR2* targets. As such, the presence of *fimR2* potentially affects the stability of these mRNAs either negatively (*fliJ*, *fliG*, *fliI*, and *fliR*) or positively (*hofQ*).

**Figure 4:**
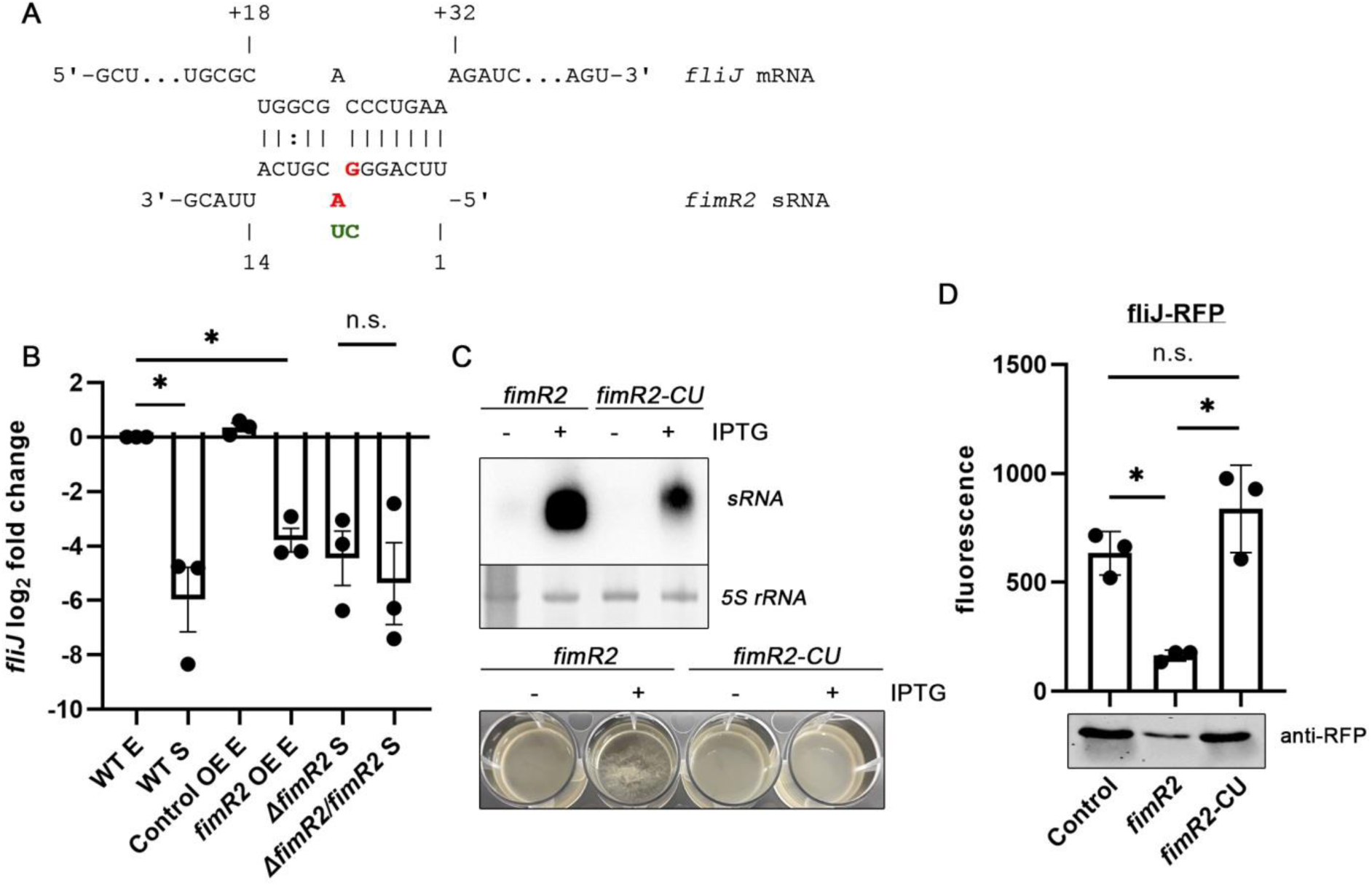
*fimR2* regulates *fliJ* mRNA at the post-transcriptional level. **A.** *fimR2*-*fliJ* predicted interaction. Mutated nucleotides are indicated in red, and their substitutions in green. **B.** RT-qPCR analysis of *fliJ* in WT, Control OE, OE, *ΔfimR2*, and *ΔfimR2/fimR2* strains. Total RNA samples from E (exponential phase) and S (stationary phase) are shown. Mean log_2_ fold change + SEM are shown from 3 biological replicates. log_2_ fold change was based on comparison with WT E samples. Unpaired two-tailed t-test with Welch’s correction was used to determine significance with n.s and * showing not significant and significant results, respectively. The p-values are 0.0373, 0.0128, and 0.6379, respectively. **C.** Northern blot analysis of *fimR2* and *fimR2*-CU (top) showing the expression of the sRNAs upon induction with IPTG. Ethidium bromide staining of *5S rRNA* is shown as a loading control. Pictures of bacterial cultures from the same conditions (bottom) showing the aggregation upon *fimR2* overexpression. **D.** Mean + SD of fluorescence of the fliJ-RFP fusion protein following control, *fimR2*, or *fimR2-CU* overexpression (top) and RFP western-blot of the same samples (bottom). Three biological replicates were used for these experiments and background fluorescence from individual sRNA overexpression strains were subtracted from experimental values. Unpaired two-tailed t-test with Welch’s correction was used to determine significance with n.s. and * showing not significant and significant results, respectively. The p-values are as follows: 0.0112 0.0268 and 0.2153.

Next, we evaluated if *fimR2* regulates these targets through direct base-pairing at the post-transcriptional level. For this purpose, the first 30 nt of *fliJ,* including the predicted target site, were fused to red fluorescent protein (RFP) and fluorescence was monitored following in the absence or presence of ectopically expressed *fimR2* in the *ΔfimR2* strain. As additional specificity control, *fimR2*-CU was tested, which is a stably expressed *fimR2* mutant defective in triggering biofilm formation (**Figure 4C**). Reporter assays showed that *fimR2* overexpression downregulated *fliJ-RFP* expression by almost 80% (**Figure 4D**). As *fimR2* overexpression caused cell aggregation and biofilm formation (**Figure 2A**), fluorescence measurements were additionally validated by western blotting (**Figure 4D**), and these experiments fully confirmed the fluorescence data.

Taken together, these findings showed that *fimR2* acts as a *trans*-acting sRNA, inhibiting the expression of *fliJ* and potentially other transcripts involved in motility (*fliG*, *fliI,* and *fliR*), thereby eliciting biofilm formation. Concomitantly, *fimR2* promotes the expression of *hofQ,* providing means for the stressed bacterium to import foreign DNA for catabolic reactions.

### *fimR2* interacts with CsrA

Bacterial sRNAs are usually stabilized by RNA chaperones, such as Hfq or ProQ, for further assisting in sRNA-target interactions (Hör et al., 2020). We thus evaluated the dependence of *fimR2* on either chaperone by monitoring its stability following *hfq* and *proQ* deletions. Northern blot analysis showed that *fimR2* abundance was unaffected in stationary phase in the absence of either chaperones (**Figure S5A**). Furthermore, *fimR2* was also not recovered in RIL-seq analyses of both chaperones (Melamed et al., 2020). Taken together, these observations suggested that, unlike most *trans*-encoded sRNAs, *fimR2* acts independently of both Hfq and ProQ.

Another prominent bacterial chaperone protein is the translational regulator CsrA that is involved in a plethora of cellular activities (Pourciau et al., 2020). Interestingly, a recent CLIP-seq analysis revealed that CsrA interacts with both the *fimA* sequence and the intergenic *fimA-fimI* region (Potts et al., 2017). As the latter is a short region composed predominantly of the *fimR2* sequence, we hypothesized that CsrA is interacting with the sRNA. CsrA favorably binds GGA motifs, with a preference to those located in secondary structures (Potts et al., 2017). The *fimR2* sequence contains two GGA motifs with the first located in the single-stranded region, and the second in the loop structure (**Figure S5B**), suggesting that the sRNA could use both motifs to interact with a CsrA dimer. To validate this potential interaction, we expressed and co-immunoprecipitated his-tagged CsrA in stationary phase and revealed a striking enrichment of *fimR2* in the eluted fractions (**Figure 5A**). The CsrA-interacting sRNA *CsrB* (Liu et al., 1997) was also recovered in the same fractions (**Figure 5A**). However, full-length *tRNA^Gly^* and its 5’-tRNA fragment, a stationary-phase enriched sRNA that we previously recovered (Raad et al., 2021), were both largely depleted from these fractions (**Figure 5A**) thus highlighting the specificity of the experimental system. Taken together, these pulldown data confirmed that *fimR2* and CsrA interact *in vivo*.

**Figure 5:**
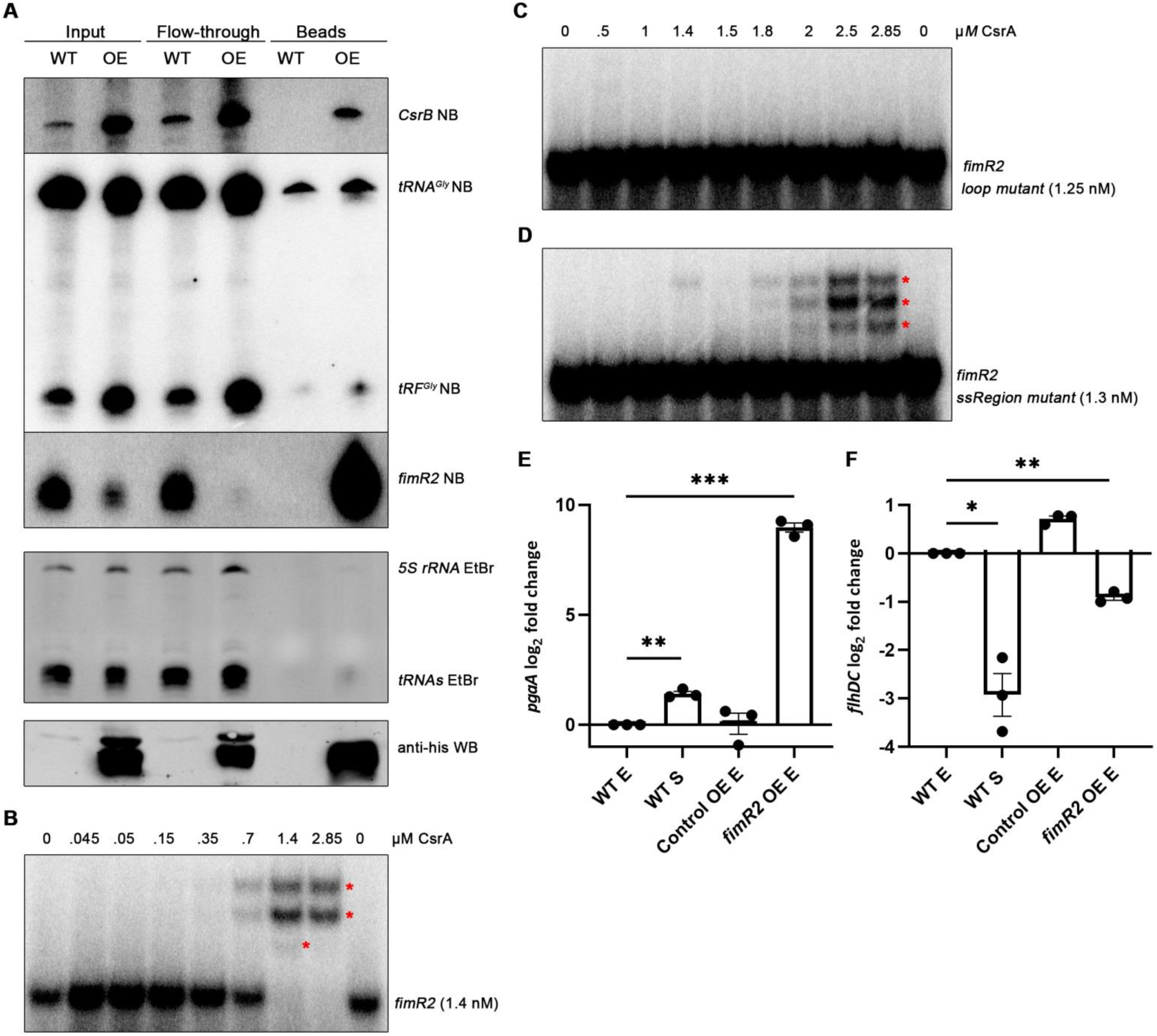
*fimR2* sequesters CsrA from its targets. **A.** Northern blot of *CsrB, 5’-tRF^Gly^,* and *fimR2* (top) from CsrA-his_10_ CoIP fractions (top) and western blot of CsrA-his_10_ from the same samples (bottom). Samples shown are taken from the stationary phase of growth of WT and OE (CsrA-his_10_ overexpression) strains. Ethidium bromide staining of *5S rRNA* and *tRNAs* are shown as loading controls. **B.** EMSA of radioactively labelled *fimR2* with increasing concentrations of purified CsrA-his_10_. **C.** EMSA of radioactively labelled *fimR2* loop mutant with increasing concentrations of purified CsrA-his_10_. **D.** EMSA of radioactively labelled *fimR2* ssRegion mutant with increasing concentrations of purified CsrA-his_10_. Upshifts in B-D are marked with red asterisks. **E.** RT-qPCR analysis of *pgaA* and **F.** *flhDC* expression from WT, Control OE, and *fimR2* OE. Samples from E (exponential phase) and S (stationary phase) are shown. Mean log_2_ fold change + SEM are shown for both transcripts from 3 biological replicates. log_2_ fold change was based on comparison with exponential phase samples. Unpaired two-tailed t-test with Welch’s correction was used to determine significance with * showing significant results. The p-values are **E.** 0.0058 and 0.005, **F.** 0.0221 and 0.0046.

To corroborate these findings, we carried out electrophoretic mobility shift assays (EMSA) with *in vitro* transcribed and 5’-end labelled *fimR2* and purified CsrA. These EMSA experiments showed that *fimR2* upshifts with increasing concentrations of purified CsrA (**Figure 5B**). Mutations of the GGA motif in the *fimR2* stem-loop region, but not in its single-stranded region (ssRegion), abolished association with CsrA (**Figure 5C, D**). Taken together, these experiments suggested that CsrA *binds fimR2* at the stem-loop region.

We next wondered what purpose this interaction is fulfilling. As *fimR2* did not associate with Hfq or ProQ (**Figure S5A**), we hypothesized that CsrA may be stabilizing the sRNA. As the deletion of CsrA is detrimental to bacterial growth in LB (Timmermans and Melderen, 2009), we thus altered cellular CsrA availability to test *fimR2* dependence on the chaperone. The CsrA translational regulator is sequestered from its target through its interaction with sRNAs *CsrB* and *CsrC* as both sRNA are heavily folded and contain multiple GGA motifs that entrap CsrA (Liu et al., 1997, Weilbacher et al., 2003). These sRNAs are degraded by RNAse E through the adaptor activity of CsrD (Suzuki et al., 2006). As such, deletion of CsrD upregulated *CsrB* and *CsrC* expression (**Figure S5C**) and thus limited the pool of available free CsrA dimers in the cell. Nonetheless, *fimR2* expression was not perturbed upon CsrD deletion **(Figure S5C**), suggesting that the sRNA interaction with CsrA is not critical for its stability. Subsequently, deletion of either *CsrB* or *CsrC* sRNAs or both would release free CsrA dimers in the cell, however neither deletion significantly boosted *fimR2* expression (**Figure S5D**). Furthermore, the addition of purified CsrA to *in vitro* cleavage reactions did not enhance the RNase E-dependent processing of the *fimR2* precursor (**Figure 1D**). We thus concluded that despite the *fimR2*/CsrA interaction formation, CsrA does not stabilize *fimR2 in vivo* and thus this interaction must be fulfilling another distinct function.

### *fimR2* sequesters CsrA from its targets

The interaction of CsrA with sRNA partners have largely been associated with antagonizing mechanisms with the sRNAs sequestrating the translational regulator from its targets (Liu et al., 1997, Weilbacher et al., 2003, Jørgensen et al., 2013) or the sequestration of the sRNA from its targets by the chaperone (Potts et al., 2017). We thus considered the hypothesis that *fimR2* is sequestering CsrA from its targets, rather than relying on the chaperone for finding and binding target mRNAs. Interestingly, SEM showed depletion of T1P in conditions giving rise to prominent *fimR2* expression (**Figure 3B, D, E**). Under similar conditions, RT-qPCR analysis revealed significant downregulation of *fimA* (**Figure S6A**) and other individual members of the *fim* polycistron (**Figure S6B-G**). As the sequence of the *fimAICDFGH* operon and its 5’-UTR do not possess predicted base-pairing sites with *fimR2, and fimR2* ectopic overexpression dramatically downregulates T1P (**Figures 3D** **and S6A-G**), we proposed that *fimR2* could regulate the expression of these transcripts indirectly. CsrA has been shown to stabilize *fimAICDFGH* by interaction with *fimS* (Mitra et al., 2013). In this manner, the interaction of *fimR2* with CsrA could be antagonizing this stabilization of the *fim* transcript, suggesting that *fimR2* is sequestering CsrA and antagonizing its activity.

Prominent morphological features seen following *fimR2* overexpression on SEM include increased PGA synthesis and a dramatic loss of flagellation (**Figure 3D**). The secretion of PGA is done by the coordinated actions of the members of the *pgaABCD* operon (Itoh et al., 2008). CsrA downregulates the expression of this transcript by allowing premature rho-dependent transcription termination (Figueroa-Bossi et al., 2014). RT-qPCR analysis showed dramatic upregulation of *pgaA* upon *fimR2* overexpression (**Figure 5E**), suggesting that binding of *fimR2* to CsrA prevented the latter from downregulating *pgaA*. In parallel, CsrA promotes flagellar motility by stabilizing the *flhDC* transcript and shielding it against RNase E cleavage (Yakhnin et al., 2013). RT-qPCR analysis of *flhDC* shows repression upon *fimR2* overexpression (**Figure 5F**) presenting the opposite scenario of the regulation by CsrA. These two experiments present so far unknown examples of CsrA antagonisms by *fimR2* and suggest that *fimR2* sequesters CsrA away from its targets in similar fashions as reported for *CsrB*, *CsrC*, and other sRNAs (Pourciau et al., 2020). Furthermore, this sequestration model is compatible with the observed prominent biofilm formation upon *fimR2* overexpression (**Figure 2A-D**), resembling those seen upon CsrA deletion (Jackson et al., 2002)

### fimR2 promotes S. enterica invasion

As the sequence of the sRNA is conserved in many *Escherichia* and *Shigella* species (**Figure S1B**), we wondered if *fimR2* is present in pathogens from the *Salmonella* genus. In a recent study, RNase E was shown to cleave near the *fimA* stop codon in *S. enterica* serovar Typhimurium (SL1344, SB300) (Chao et al., 2017) generating a small transcript whose sequence is similar to that of *E.coli fimR2* (**Figure 6A**). We refer to this putative sRNA as *fimR2S* to denote it as the *fimR2* variant in *Salmonella*. Northern blot analyses demonstrated the expression of a ∼30-nt RNA in late stationary phase in the assayed *Salmonella* strains (**Figure 6B**). This finding and the reported RNase E cleavage site within the *fimA* transcript suggested a similar phase-dependent cleavage occurring in stationary phase and releasing *fimR2S* from the *fimA* transcript. *S. enterica* often express T1P under laboratory growth conditions and under static conditions (Hansmeier et al., 2017). This suggests that the lower abundance seen for *fimR2S* as compared to *E. coli* is due to the continuous T1P expression in these *Salmonella* strains.

**Figure 6:**
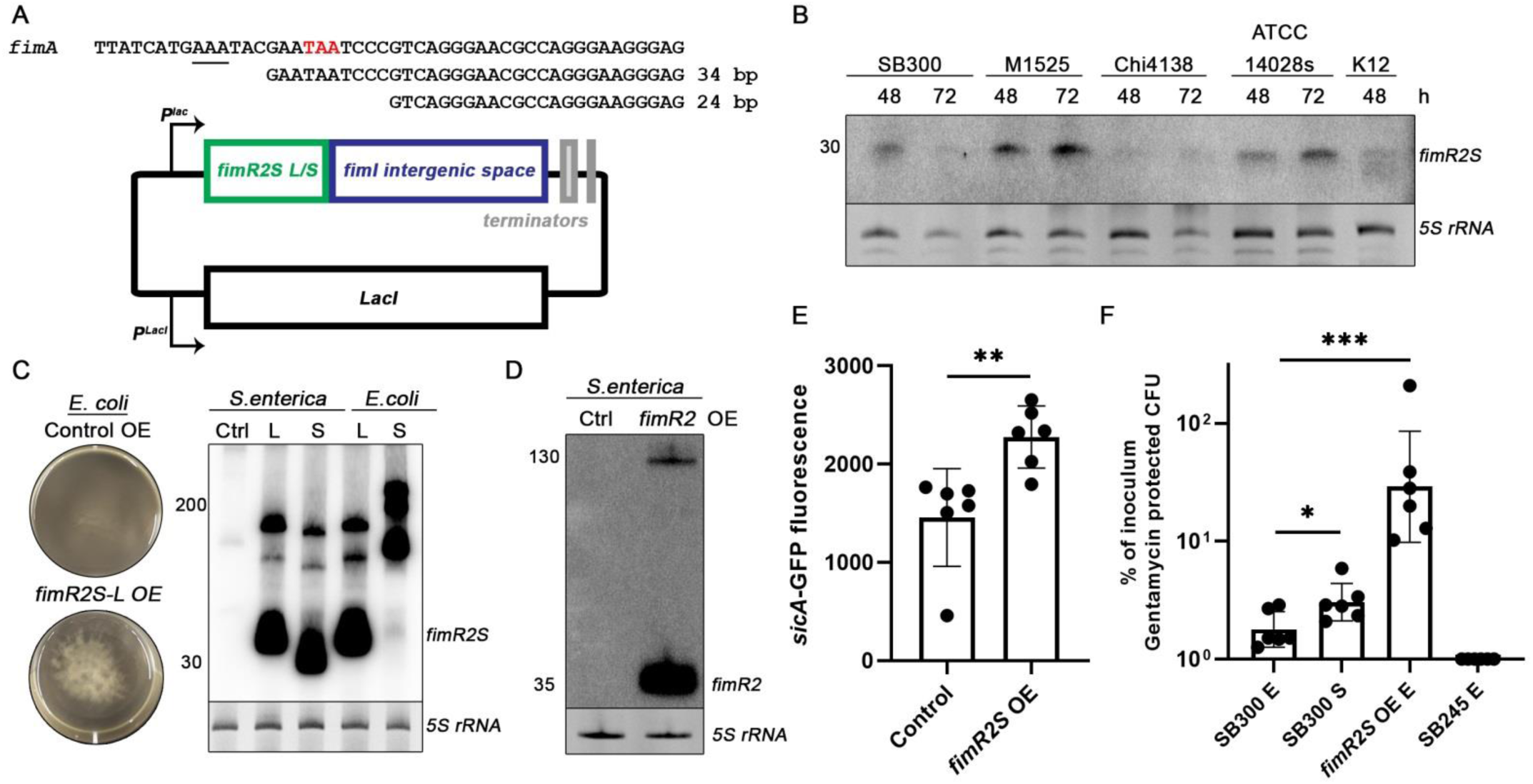
Salmonella *fimR2S* is involved in infection. **A.** 5’-RACE results mapped to *S. enterica fimA* sequence (top) and *fimR2S* expression plasmid (bottom). *fimA* stop codon is indicated in red. The predicted sequence for RNase E cleavage based on analysis from Chao et al. (2017) is underlined. Sizes of recovered transcripts are indicated in bp (base pairs). **B.** Northern blot analysis of *fimR2S* in four *Salmonella* strains and the *E. coli* K12 strain. Total RNA samples from different time points of bacterial growth, 48 and 72 h (hours), are shown. Ethidium bromide staining of *5S rRNA* is shown as a loading control**. C.** Pictures of Control and *fimR2S-L* OE in *E. coli* (left) and northern blot analysis of *fimR2S* expression in *S. enterica* and *E. coli* upon overexpression of *fimR2S-L* and *fimR2S-S*, the long and short *fimR2S* variants, respectively. Ctrl designate overexpression of the control. Ethidium bromide staining of *5S rRNA* is shown as a loading control. The size of the different RNA molecules is indicated on the left. **D.** Northern blot analysis of *fimR2* expression in *S. enterica* under control and *fimR2* OE conditions. Ethidium bromide staining of *5S rRNA* is shown as a loading control**. E.** Mean + SD of *sicA*-GFP fluorescence without (Control) or with *fimR2S* OE from six biological replicates. Unpaired two-tailed t-test with Welch’s correction was used to determine significance with ** showing significant results and p-value of 0.0086 as compared to the control samples. **F.** Mean + SD of percentage of inoculum protected from gentamicin treatment following infection of HeLa cells with an initial inoculum of SL1344 (SB300) from E (exponential phase) and S (stationary phase), and *fimR2S* overexpression strains, from six replicate infections. SB245 strain is used as a negative control for invasion. Calculations were done following counting of colony forming units (CFU) of protected cells and initial inocula. Unpaired two-tailed t-test with Welch’s correction was used to determine significance with * showing significant results. The p-values are, in order, 0.0280 and 0.0010.

To understand the potential functions of *fimR2S*, we overexpressed it in both *S. enterica* and *E. coli* followed by tracking the phenotypic changes. To this end, we conducted 5’- and 3’-RACE experiments to map the exact ends of this sRNA. Only 5’-RACE experiments were successful (**Figure 6A**) and showed a lower abundance, longer transcript (*fimR2S-L*) that retains part of the *fimA* coding sequence, and a shorter more abundant form (*fimR2S-S*) solely generated from the *fimA-fimI* intergenic space. We thus cloned both forms into an expression plasmid (**Figure 6A**). To generate the canonical 3’-end of the transcript, we included a portion of the *fimA-fimI* intergenic space downstream of the two variants, prompting RNAse E to generate the canonical 3’-end of *fimR2S* or for its potential secondary structure to terminate its expression. Strikingly, overexpression of *fimR2S-L*, but not *fimR2S-S* in *E. coli* caused biofilm formation (**Figure 6C**) as only the longer variant was processed properly. Overexpression of either *fimR2S* variants in *S. enterica* did not promote biofilm formation (data not shown). Similarly, the overexpression of the *E. coli fimR2* in *S. enterica* (**Figure 6D**) did not promote biofilm formation. Taken together, these experiments suggested that *Salmonella fimR2S* mimics *fimR2* functions in *E. coli* but the sRNA is not functional in *S. enterica*. Alternatively, *fimR2S* may be operating in *S. enterica* differently than in *E. coli*.

*S. enterica* infectivity largely relies on the secretion of several effectors by the T3SS (Galán, 2021). Considering the effects of *fimR2* on biofilm formation (**Figure 2D**) and outer membrane architecture (**Figure 3D**) in *E. coli,* we hypothesized that the sRNA is involved in virulence and used *fimR2S* in *S. enterica* to evaluate this possibility. SicA is a critical T3SS chaperone that promotes the expression of invasion mediating effector proteins (Tucker and Galán, 2000). Significantly, *fimR2S* overexpression induced sicA-GFP fusion protein (Sturm et al., 2011) in a reporter assay (**Figure 6E**), suggesting that the *Salmonella* sRNA is functional in promoting T3SS secretion. Next, we hypothesized that the sRNA could regulate infectivity as it positively regulates T3SS. To test this hypothesis, we infected HeLa cells with wild-type *S. enterica* from both exponential and stationary phase and with the *fimR2S* overexpression strain (Pfister et al., 2020). Using the *SB245* strain, which is invasion deficient as a negative control, we quantified the intracellular bacterial load following treatment with gentamicin, which is membrane impermeable and consequently kills non-invaded extracellular bacteria selectively. According to this invasion assay, *fimR2S* overexpressing strains were markedly more invasive than either wild-type strains (**Figure 6F**), suggesting that *fimR2S* promotes invasion and internalization of bacterial cells into human cells.

## Discussion

In this study we identified and functionally characterized *fimR2*, an abundant 35 nucleotide-long sRNA specifically expressed in stationary phase of bacterial growth (**Figure S1A**). To this end, we ectopically overexpressed this sRNA during exponential phase to investigate its potential regulatory effects outside its canonical growth phase context. While the overexpression in exponential phase produced much more sRNA than is normally present in stationary phase (**Figure 2C**), the observed phenotype (namely regulation of fimbrial and flagellar mRNAs, modulation of outer membrane architecture, and induction of biofilm formation) was comparable to the effects in stationary phase. We report the first functional characterization of this sRNA, and with the generated knowledge we confidently ascribe a master regulatory status to *fimR2*, an RNase-E dependent sRNA which single-handedly and without the assistance of known RNA chaperones, coordinates motility, biofilm formation, stationary phase biology, and even virulence (**Figure 7**).

**Figure 7:**
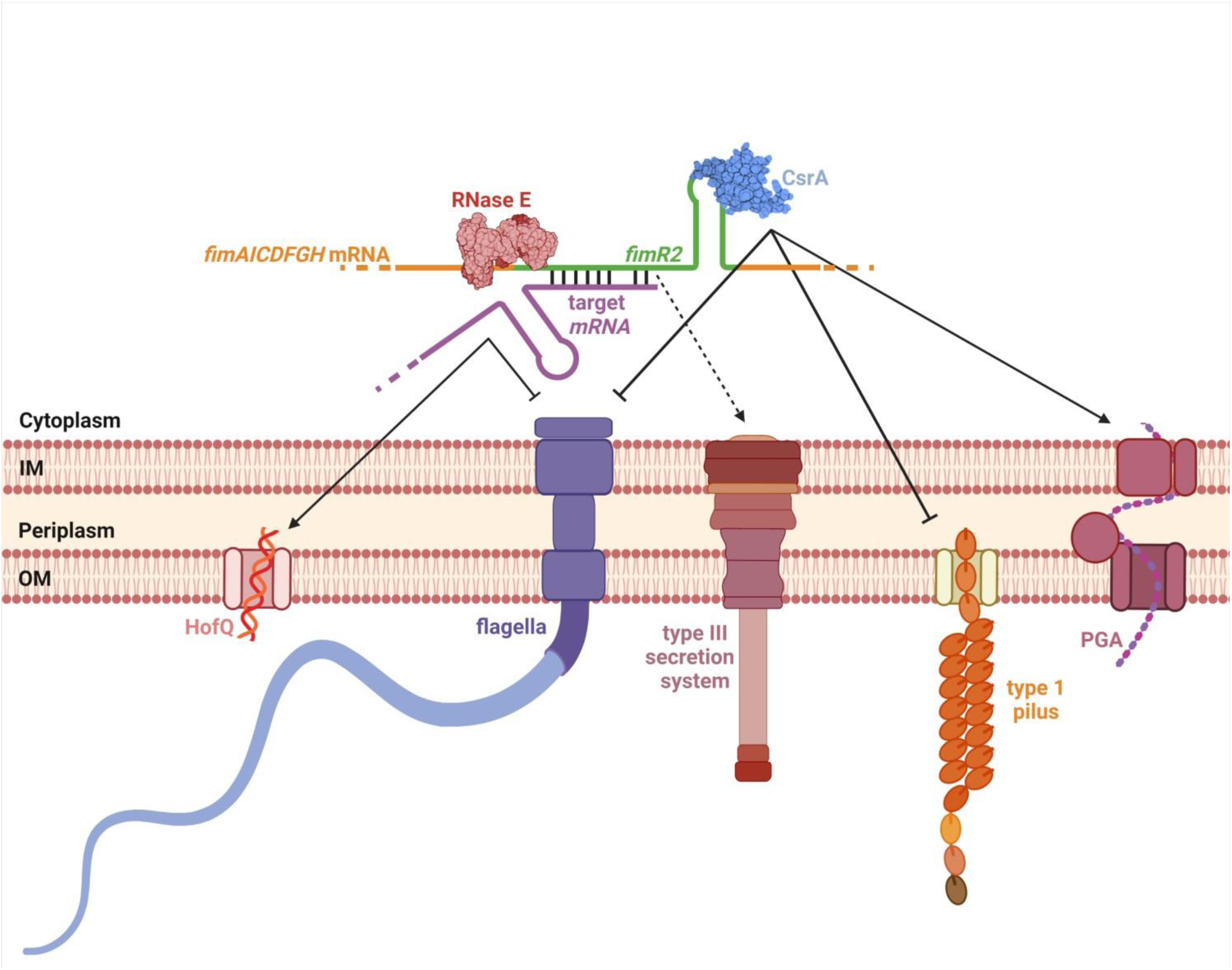
Model of *fimR2* function. *fimR2* sRNA is processed from the *fimAICDFGH* transcript in stationary phase by RNase E. The sRNA (green) interacts with the translational regulator CsrA and antagonizes its effects, upregulating PGA synthesis, and downregulating type 1 pilus and flagellar synthesis. *fimR2* upregulates type III secretion likely through the sequestration of CsrA. In parallel, *fimR2* regulates several transcripts through direct base-pairing, inhibiting flagellar synthesis and upregulating HofQ-mediated import of extracellular DNA for use in catabolic reactions. The crystal structure of RNase E catalytic domain (2C4R) and the NMR structure of CsrA (1Y00) were used (Callaghan et al., 2005, Gutiérrez et al., 2005). This figure was created with www.biorender.com.

### *fimR2* orchestrates biofilm formation through multitasking

*fimR2* overexpression instructed bacteria to a rapid transition to a sessile lifestyle (**Figure 2A-D**). *fimR2* orchestrates this complex shift in multiple ways: the sRNA upregulated *hofq* mRNA involved in stationary phase biology (**Figure S4D**) and downregulated several targets coding for flagellar protein synthesis (**Figures 4B, D** **and S4A-C**), explaining the shift from planktonic to sedentary growth stages. In this context, *fimR2* appears to act in a similar fashion to other bacterial *trans*-encoded sRNAs, regulating target mRNAs at the posttranscriptional levels (Nitzan et al., 2017). As the mRNA levels of several predicted targets directly responds to *fimR2* varying levels, *fimR2* is likely recruiting RNases to degrade certain targets (*fliJ, fliG, fliI,* and *fliR*) while stabilizing others (*hofQ*).

In a separate mode of action, *fimR2* sequesters CsrA, a known translational regulator of the Type III secretion system, from its target genes, which are involved in bacterial motility and biofilm formation. CsrA affects the levels of *flhDC* (motility) and *pgaABCD* (biofilm formation) mRNAs, respectively, by shielding the former from RNase degradation and allowing Rho access to the transcript of the latter (Yakhnin et al., 2013, Figueroa-Bossi et al., 2014). By downregulating *flhDC* expression (**Figure 5F**) and upregulating *pgaA* expression (**Figure 5E**), *fimR2* appears to antagonize these specific CsrA functions. In a similar manner, *fimR2* ectopic overexpression downregulated *fimA* expression (**Figure S6A**). While the sRNA is processed from the complete *fimAICDFGH* operon mRNA in stationary phase (**Figure 1B, C)** and through this mode, the parental transcript is downregulated, *fimR2* likely tightens the inhibition on the *fim* operon by competing for CsrA binding (Mitra et al., 2013).

Finally, *fimR2* overexpression caused the upregulation of *rpoS*, the stationary phase-dependent RNA polymerase sigma factor (Raad et al., 2021). While this inductive effect is likely indirect, as we do not observe a putative base-pairing site for *fimR2* within the *rpoS* mRNA sequence, it explains the propensity of bacterial cells to form biofilms (**Figure 2A**) and to adopt stationary phase-like morphology (**Figure 3B, D, F**). Future work is needed for a better understanding of the mechanism by which *fimR2* affects *rpoS* expression.

### Efficiency and sustainability in bacterial regulatory networks

The simultaneous modulation of biofilm formation, motility, stationary phase biology, and even virulence (**Figure 7**) is not a trivial task. The success of *fimR2* in these strenuous activities and its ability to navigate several branches within a complex regulatory network can be attributed to two distinct features: *fimR2* dual mode of function and its abundance. Bacterial *trans*-encoded sRNAs are efficient and potent regulatory molecules as they typically employ their single-stranded regions to base-pair and regulate various targets mRNAs at the post-transcriptional level (Nitzan et al., 2017). Indeed, the range of *fimR2* mRNA targets is very diverse (**Figure S4A-D**) and includes transcripts coding for flagellar proteins and porins, among many others. Another mode by which *fimR2* exhibits utmost efficiency is through its sequestration of the translational regulator CsrA (**Figure 5E, F**). CsrA itself is a master regulator of gene expression, governing diverse cellular processes including motility, biofilm formation, virulence, quorum sensing, metabolism, and oxidative stress (Pourciau et al., 2020). By sequestering CsrA from mediating its cellular functions, *fimR2* automatically gains regulatory access to the CsrA-specific expression network and navigates its intricate web, remodeling its fibers in an antagonistic manner. Such a dual regulatory function has also been attributed to other sRNAs such as *McaS* (Jørgensen et al., 2013) that simultaneously regulates target mRNAs and sequester CsrA. Furthermore, CsrA has been shown to interact with other sRNAs (Potts et al., 2017). With the existence of these various parallel examples, it is now more evident that bacterial species have evolved efficient and elegant regulatory modes to rapidly adapt to unpredictable environments.

*fimR2* abundance is another feature contributing to its potent effect. Indeed, the sRNA is far more abundant than its peers (*ArcZ* for example, **Figure 1B**) explaining its ability to tightly modulate various targets simultaneously. Furthermore, *fimR2* is more abundant than *CsrB* and *CsrC* (**Figure S5C, D**). While these two sRNAs possess multiple GGA motifs to entrap CsrA in their secondary structures, the single GGA motif on the highly abundant *fimR2* molecule is sufficient to efficiently sequester CsrA (**Figure 5A**). While the expression of such an abundant RNA molecule would normally entail overwhelming energy expenditure, *fimR2* biogenesis constitutes another efficient mechanism. Culminating *in vivo* and *in vitro* evidence revealed that RNase E processes *fimR2* from the parental *fim* transcript (**Figure 1B-D**). As the accumulation of this sRNA occurs in stationary phase, the processing event likely overlaps with *fimAICDFGH* mRNA turnover during the same growth phase. *fimAICDFGH* is very abundant in *E. coli* (**Figure S1A**) and in *Salmonella* (Sterzenbach et al., 2013), explaining the abundance of the resulting sRNA.

### *fimR2* is an RNase-E dependent sRNA

What signals are required for this specific *fimR2* cleavage to occur locally and temporarily are still elusive. RNase E determinants include AU-rich sequences, stem-loop structures, and transcripts with 5’-monophosphates (Updegrove et al., 2018). The *fimA* deletion strain used in this study retains a portion of the *fimA* ORF at the 3’-end, near the stop codon (Baba et al., 2006). As this knockout did not reduce *fimR2* expression (**Figure S2B**), we postulate that any sequence-specific motifs or secondary structures required for *fimR2* processing by RNase E are present in its vicinity. That a 40-nt intermediate was generated upon *in vitro* RNase E cleavage of a *fimR2* precursor (**Figure 1C, D**) suggests a role for another RNase in complete *fimR2* biogenesis. While our consideration of other RNases (**Figure S2D, E**) was not comprehensive, the depletion of *fimR2 in vivo* upon RNase E deactivation (**Figure 1B**) speaks against additional RNases being involved. Furthermore, *in vitro* cleavage assays with total cell lysates generated the same 40-nt intermediate (data not shown) suggesting the *in vitro* precursor transcript lacking crucial determinants for complete processing, such as post-transcriptional modifications or the establishment of correct three-dimensional architecture.

Finally, the secondary structure adopted by *fimR2* resembles that of an intrinsic terminator (**Figure S5B**). Indeed, a similar structure has been proposed to serve as an intrinsic terminator for *fimA* expression (Klemm, 1984). As such, a single processing event occurring at the 3’-end of the *fimA* transcript, and thus at the 5’-end of the *fimR2* sRNA, would have to take place to release the sRNA from the parental transcript for final processing by RNase E. This scenario can explain the observed proper transcription termination in the *fimR2* and *fimR2S* overexpression experiments **(Figures S3B and 6C, D),** and the appearance of the 40-nt intermediate (**Figure 1C, D**).

### *fimR2* does not associate with Hfq or ProQ

*fimR2* is a functional sRNA possessing regulatory roles independently of the known RNA chaperones Hfq and ProQ (**Figure S5A**). This finding is surprising as most *trans*-encoded regulatory sRNAs in bacterial species require a chaperone to maintain their stabilities or to anneal to their target mRNAs (Hör et al., 2020). In variance, *fimR2* interacts with the bacterial translational regulator CsrA (**Figure 5A**). CsrA seems to bind *fimR2* via its predicted stem-loop structure (**Figure 5C**) which is in agreement with the reported preference of CsrA to bind GGA motifs in stem-loop structures (Potts et al., 2017). The interaction with CsrA does not seem to stabilize the sRNA as depletion of the pool of available CsrA molecules did not perturb *fimR2* abundance (**Figure S5C**). The opposite is also true as the increase of CsrA molecules did not enhance *fimR2* stability (**Figure S5D**). Forming yet another contrasting phenomenon to other sRNAs, *fimR2* processing by RNase E did not require a chaperone (**Figure 1C**) nor did it depend on CsrA (**Figure 1D**).

Mutagenesis of *fimR2* abolished its regulatory functions (**Figures S3C, D, and 4C, D**). However, only the mutations targeting the predicted secondary structure impaired *fimR2* stability (**Figure S3B**). As *fimR2* does not seem to associate with any known RNA chaperon, this secondary structure may be in fact the only factor required for the sRNA stability. In support of this interpretation, unfolded *fimR2* was rapidly degraded upon incubation with total cell lysates (data not shown), suggesting that the predicted stem-loop is indeed critical for stability. However, it is possible that *fimR2* associates with a so far uncharacterized RNA chaperon *in vivo*, a hypothesis compatible with the observed upshift patterns seen in crosslinking experiments (**Figure S5E**).

### *fimR2* contributes to virulence

This work illustrates several modes by which *fimR2* promotes stationary phase biology (**Figure S4D**), biofilm formation (**Figures 2B-D** **and 5E**) and suppresses bacterial motility (**Figures 3D, 5F, and S4A-C**). All three processes have been shown to be important for bacterial virulence. Bacteria display increased expression of virulence genes during stationary phase (Navarro Llorens et al., 2010). Pathogens also use their flagella to swim towards sites of infection (Josenhans and Suerbaum, 2002). They form biofilms as a form of resistance against antimicrobial treatments and persistence within hosts (Naziri et al., 2021). Furthermore, bacterial biofilms on medical equipment significantly contribute to infections (Wang et al., 2018). In this context, we noted that biofilms resulting from *fimR2* overexpression (**Figure 2B**) or *fimE* deletion (**Figure S2F, G**) were extremely resistant to mechanical stress (centrifugation and pipetting during SEM preparation) as well as chemical stress through ethanol and SDS treatment. With the contribution of *fimR2* to these processes, it is thus logical to posit that *fimR2* likely contributes to disease establishment. In fact, CopraRNA analysis predicted several targets to be involved in lipopolysaccharide (LPS) synthesis, suggesting that *fimR2* could modulate LPS synthesis and display, a prerequisite for disease establishment.

To investigate *fimR2* role in virulence, we use *fimR2S*, the *fimR2* homologue of *S. enterica* as a proxy. While *fimR2S* overexpression in *S. enterica* did not cause detectable biofilm formation (data not shown), it promoted biofilm formation in *E. coli* (**Figure 6C**) suggesting that *fimR2S* is functional and can substitute for *fimR2* in *E. coli*. Though *E. coli fimR2* overexpression in *S. enterica* was successful (**Figure 6D**), it did not contribute to biofilm formation either. It is tempting to assume that *fimR2S* operates in *S. enterica* in different modes although it is generated from the same genomic context and likely through similar RNase E cleavage. Because of the involvement of *fimR2* in biofilm formation (**Figure 2A-D**), we hypothesized that the sRNA or its homologue in *S. enterica*, *fimR2S*, would be involved in virulence. Indeed, *fimR2S* overexpression in *S. enterica* upregulates sicA expression (**Figure 6E**), paralleling the upregulation of this T3SS-chaperone upon CsrA mutation (Hung et al., 2019). And indeed, *fimR2S* overexpressing strains were clearly more infectious of HeLa cells than their wild-type counterparts (**Figure 6F**). These two experiments suggested that *fimR2S* can promote *S. enterica* virulence via a currently unknown mechanism. Like *fimR2*, *fimR2S* also retains two GGA motifs (**Figure 6A**) that could be employed to interact with CsrA. Furthermore, the 5’-UTR of the *Salmonella fim* transcript has been reported previously to interact with CsrA (Sterzenbach et al., 2013). Based on these similarities, we hypothesized that in *S. enterica*, *fimR2S* can function to sequester CsrA from its targets in a comparable manner as in *E. coli*. While we did not investigate the association between CsrA and the sRNA in this species, *fimR2S* promotion of virulence suggests potential sequestration of CsrA. As CsrA plays critical roles in regulating T3SS effectors (Altier et al., 2000, Lou et al., 2019) and thus disease establishment, it is thus tempting to assume that the *fimR2S*-dependent induction of *sicA* occurs through the sequestration of CsrA. More experimental work is needed to validate these hypotheses.

Finally, the growth phase dependent expression of *fimR2* in the *E. coli* clinical isolates *ESBL*, *VTEC*, and *UPEC* strains (**Figure S1C, D**) suggests a role for the sRNA in these strains. K12 and UPEC strains show similar predicted targets for *fimR2* regulation suggesting that the sRNA could potentially play similar regulatory functions in the pathogenic strain. Furthermore, the dependence of UPEC on the T1P suggests that *fimR2* might play important roles in dictating the expression of the conserved *fimAICDFGH* operon. The expression of *fimR2* in pathogenic *E. coli* then can remodel the outer membrane architecture by inhibiting T1P expression and promoting that of other adhesins. In this context, the use of *UPEC* as a model to study *fimR2* expression during invasion can provide better understanding of the function of the sRNA in disease.

In summary, this study revealed a so far functionally uncharacterized 35 nucleotides long non-coding RNA, named *fimR2*, to serve as post-transcriptional regulator in *E. coli* and *Salmonella*. *fimR2* is a stationary phase-specific sRNA that modulates several target mRNAs with the result of increased biofilm formation, reduced motility and enhanced infectivity. *fimR2* might play an even more intricate role as global regulator on the population level during infections, as it has been demonstrated to be one of the most abundant secreted RNA molecules in outer membrane vesicles (Ghosal et al., 2015). Therefore our study can be regarded as steppingstone for future dedicated work on *fimR2* for elucidating even more elaborate mechanisms by which this tiny RNA molecule weaves its regulatory network.

## Supporting information

Strains and plasmids

Oligos

## Acknowledgments

We would like to thank Alexander Mankin (UIC, Chicago) for providing KEIO strains, Eric Massé (University of Sherbrooke) for sharing the *rne3071^-ts^* strain, Ben Luisi (University of Cambridge) for sharing active RNE-NTD fractions, and Markus Hilty (IFIK, Bern) for providing *E. coli* human and pathogenic strains. We would also like to thank Andrew Hemphill (VetSuisse, Bern) for providing expertise and reagents for SEM, and Beatrice Frey (DCBP, Bern) for assisting with the Zeiss Microscope. This work was funded by the Swiss National Science Foundation (grant 310030-197515 to N.P). Additional support from the NCCR “RNA & Disease” (to N.P.) and the NCCR “Microbiomes” (to S. H.) funded by the Swiss National Science Foundation is acknowledged. D.T. was a recipient of a Helmut-Horten-Foundation grant.

## Author Contribution

Conceptualization, N.R. and N.P.; Methodology, N.R., T.D. and S.H.; Formal Analysis, N.R.; Investigation, N.R.; Resources, N.R.; N.P.; T.D. and S.H.; Writing – Original Draft, N.R.; Writing – Review and Editing, N.P., N.R., and S.H.; Supervision, N.P.; Funding Acquisition, N.P.

## STAR Methods

### RESOURCE AVAILABILITY

#### Lead contact

Further information and requests for resources and reagents should be directed to and will be fulfilled by Norbert Polacek

#### Materials availability

The strains and plasmids generated in this study will be provided upon request.

#### Data and code availability

This paper does not report any original code.

### EXPERIMENTAL MODEL AND SUBJECT DETAILS

Strains are included in Table S1 and their conditions of growth are described below.

#### Key Resource Table

| REAGENT or RESOURCE | SOURCE | IDENTIFIER |
| --- | --- | --- |
| **Antibodies** |  |  |
| 6*His, His-Tag mouse monoclonal antibody | Proteintech | Catalogue number: 66005-1-Ig |
| Anti-RFP mouse monoclonal antibody | Rockland | Number: 200-301-379S |
| IRDye 800CW Goat anti-mouse IgG secondary antibody | Licor | AB_621842 |
| **Bacterial and virus strains** |  |  |
| See Table S1 |  |  |
| **Critical commercial assays** |  |  |
| Wizard<sup>®</sup> SV Gel and PCR Clean-Up | Promega | A9282 |
| Wizard<sup>®</sup> Plus SV Minipreps DNA Purification System | Promega | A1460 |
| SuperScript IV Reverse Transcriptase | Thermo Fisher Scientific | Catalogue number: 18090050 |
| FirstChoice<sup>™</sup> RLM-RACE Kit | Invitrogen | AM1700M |
| GoTaq qPCR Mastermix | Promega | A6002 |
| pGEM-T-Easy Vector System | Promega | A1360 |
| Pierce BCA protein assay kit | Thermo Fisher Scientific | 23225 |
| **Experimental models: Cell lines** |  |  |
| HeLa (Kyoto) | S. Hapflmeier, IFIK, University of Bern |  |
| **Oligonucleotides** |  |  |
| See Table S2 |  |  |
| **Recombinant DNA** |  |  |
| See Table S1 |  |  |

#### Methods

##### Bacterial strains and growth conditions

*E. coli* K12 strain and its derivatives (**Table S1**) were grown at 37°C with shaking at 220 rpm in LB (10 g tryptone, 10 g NaCl, and 5 g yeast extract per L). Exponential phase samples were taken at OD_600_ = 0.4 or at indicated time points after reaching this OD_600_. Stationary phase samples were taken 24, 48, or 72 h after initial inoculation. Induction with IPTG (Isopropyl β-d-1-thiogalactopyranoside) or tetracycline was done at OD_600_ = 0.4. All media were supplemented with kanamycin (25 μg /mL), ampicillin (100 μg /mL), streptomycin (100 μg/mL), IPTG (1mM), or tetracycline (25 nM) where indicated. For the RNase E inactivation experiment, K12 and *rne3071^-ts^* (McDowall et al., 1993) strains were grown at 30°C to stationary phase and then the cultures were transferred to 43°C. *Salmonella* strain SL1344 (Hoiseth and Stocker, 1981), M1525 (Suar et al., 2006), ATCC14028s (Fields et al., 1986), UK-1 (Dieye et al., 2009), and SB245 (Kaniga and Galan, unpublished) were grown at 37°C with shaking at 150 rpm in LB supplemented with 0.3 M NaCl. For northern blot analysis, *Salmonella* stationary phase samples were taken 48 and 72 h after initial inoculation.

##### Construction of strains and plasmids

*fimR2S-L* and *fimR2S-S* were inserted, along with their downstream intergenic space, at the transcriptional +1 site under *PlacO* control in pBbE6k-RFP (Lee et al., 2011) by MEGAWHOP cloning (Miyazaki, 2011). For this, a pair of primers (See **Table S2**) were used to create a mega primer spanning the genomic *fimR2S* sequence by PCR using Phusion DNA Polymerase (NEB). The resulting PCR product was purified with Wizard^®^ SV Gel and PCR Clean-Up (Promega) and used in excess to insert the sRNAs into the pBbE6k-RFP plasmid and replace the RFP sequence with PCR. The PCR reaction was digested with DpnI (New England Biolabs) for 2 h at 37°C and transformed into K12 *E. coli* competent cells. Plasmids were extracted from positive clones by Wizard^®^ Plus SV Minipreps DNA Purification System (Promega) and sequenced with HL0199 primer (Microsynth). *fimR2* mutants (CU, stem, loop, and A-U stem) were generated by one-step cloning PCR (Qi and Scholthof, 2008) using pBbE6k-*fimR2* plasmid as a template (Raad et al., 2021). PCR reactions were DpnI treated as previously mentioned, and subsequent transformation, plasmid extraction, and sequencing were carried out, as described for pBbE6k-*fimRS-L* and pBbE6k-*fimRS-S*. Using one-step cloning PCR, 18 base-pairs of *fliJ* sequence corresponding to codons 6-12 were inserted between the first and second codon of the RFP sequence in pBbA2A-RFP. In a subsequent step, 12 additional base-pairs of the *fliJ* sequence corresponding to *fliJ* codons 2-5 were inserted upstream of the previous *fliJ* insert.

CsrA was amplified from K12 genomic DNA by PCR with primer pairs containing BamH1 and NdeI restriction sites and the histidine tag. The PCR product was then digested with both restriction enzymes as described and purified by size-exclusion electrophoresis. Ligation into BamH1/NdeI-cut backbone (pBbE6k-RFP with RFP excised) was done with T4 DNA Ligase (Promega). Subsequent subcloning was done as standard. Clones were confirmed by sequencing using HL0030 primer at Microsynth.

Genomic deletion strains were created as described (Datsenko and Wanner, 2000). sRNA locus (*fimR2*, *CsrB*, and *CsrC*) and gene (*CsrD*)-specific primers (**Table S2**) were used to amplify a kanamycin cassette from pKD13 plasmid by PCR. Purified PCR product was electroporated into K12 *E. coli* strain containing the pKD46 plasmid and expressing lambda red recombinase through induction with 1 mM Arabinose. Positive clones were recovered on LB agar plates supplemented with Kanamycin. Homologous recombination was confirmed by PCR as described in the PCR section. For the sRNA-deletion strains, the chromosomally-inserted Kanamycin cassette was removed by FLP recombination using pcp20 plasmid as previously described (Cherepanov and Wackernagel, 1995). Removal of the cassette was also confirmed by replica plating on LB agar plates with and without antibiotics, and by PCR.

##### RNA extraction and northern blotting

RNA extraction was performed with hot acidic phenol as described (Luidalepp et al., 2016). For northern blot analyses, 5-10 μg of total RNA were separated on denaturing polyacrylamide gels (8% Acrylamide M-Bis, 7 M Urea, 1× TBE), and gels were run for 2 h at 250 V. RNA was transferred to a nylon membrane (Amersham Hybond-N+, GE Healthcare) using a semi-dry blotter (V20-SDB, Scie-Plas) and crosslinked to membranes using a microprocessor-controlled UV irradiation system (BLX-254, Vilber Lourmat). DNA oligonucleotides were end-labeled with [γ-^32^P]-ATP (Hartmann Analytic) using PNK (Thermo Fisher Scientific) and used for hybridization as described (Gebetsberger et al., 2012).

For agarose gel northern blot, 20 μg total RNA were resolved on a denaturing agarose gel (1.2% agarose, 0.5% formaldehyde, 1x MOPS). RNA samples were transferred to a nylon membrane by capillary action in 20X SSC overnight at room temperature. After crosslinking RNA to membrane as mentioned, the membrane was incubated in 2xSSC buffer then in hybridization buffer at 65°C for 1 h. The template for *fimA* probe was amplified by PCR from K12 genomic DNA then labelled with [α-^32^P]-dCTP (Hartmann Analytic) with the Klenow fragment (Thermo Fisher Scientific) for 1 h at 37°C. Hybridization was done overnight at 65°C. Two washed were carried out on the following day for 30 min each in wash buffer I (2x SSC, 0.1% SDS) and II (0.2x SSC, 0.1% SDS).

##### Terminator exonuclease treatment

Terminator exonuclease treatment was done according to manufacturer manual. One microgram of total RNA from stationary phase was mixed with 1 unit TEX (Lucigen) and 4 units of RNasin Plus RNase inhibitor (Promega) in buffer A and incubated at 30°C for 60 min. Treated RNA was then phenol-chloroform extracted and precipitated in 2.5 volumes of 100% ethanol and 0.3 M sodium acetate.

##### *In vitro* transcription

*fimR2* and *fimR2* precursor DNA templates were generated by PCR with primers containing the T7 promoter sequence (**Table S2**). After PCR cleanup with Wizard^®^ SV Gel and PCR Clean-Up (Promega), RNAs were transcribed by T7 polymerase as described (Erlacher et al., 2011). For crosslinking experiments, transcription was carried out using 4-thio-uridine (Jena Bioscience). All transcripts were purified by gel filtration with G-25 sephadex (Sigma). For 5’-end labelling for EMSA experiments and crosslinking experiments, *in vitro* transcribed *fimR2* RNAs were dephosphorylated using CIP (New England Biolabs) at 37°C for 30 min. RNA was purified by phenol-chloroform extraction and precipitation in ethanol. RNA was then further purified by size-exclusion electrophoresis and 5’-end labelled with PNK (Thermo Fisher Scientific).

##### RNase E *in vitro* cleavage assay

The purified amino-terminal domain of RNase E was provided by B. F. Luisi (University of Cambridge). The following assay was carried out as described (Updegrove et al., 2018) in with some modifications. Briefly, *in vitro* transcribed *fimR2* precursor was treated with RppH (New England Biolabs) in NEB Buffer 2.0 to remove a pyrophosphate from the triphosphorylated 5’-end of the RNA. The reaction was carried out at 37°C for 30 min then purified with a phenol-chloroform extraction. For *in vitro* cleavage, RNA was heat-denatured at 95°C for 1 min then incubated at room temperature for 10 min. Afterwards, 300 mM RNA was mixed with RNase E buffer (25 mM Tris pH = 7.5, 50 mM NaCl, 50 mM KCl, 10 mM MgCl_2_, 1 mM DTT), 4 units of RNasin Plus RNase inhibitor (Promega), and with or without 300 mM CsrA-His_10._ The reactions were incubated at 30°C for 10 min then RNase E was added to a final concentration of 300 nM (+), 600nM (++), or 900 nM (+++). The reactions were then incubated at 30°C for an additional 30 min then phenol-chloroform extracted. RNA samples were precipitated at -20°C overnight in 2.5 volumes 100% ethanol, 0.3 M sodium acetate, and glycoblue (Thermo Fisher Scientific). Precipitated RNA was resuspended in 2x RNA loading dye, resolved on denaturing polyacrylamide gels, and analyzed by northern blotting.

##### Co-immunoprecipitation and protein purification

CsrA-His_10_ was overexpressed in exponential phase using IPTG. For co-immunoprecipitation experiments, 50 ml cultures were harvested on the following day (24 h, stationary phase) from K12 and K12/pBbE6k-CsrA-His_10_. Bacterial pellets were lysed in lysis buffer (50 mM NaH_2_PO_4_, 300 mM NaCl, 10 mM imidazole at pH = 8.0), 1 mg/mL lysozyme, and 10 mM VRC (New England Biolabs) for 30 min on ice. After sonication (6 rounds of 10 s, with 10 s pausing on ice in between), lysates were passed through a narrow-gauge needle to disrupt genomic DNA. Cell lysates were cleared through centrifugation at 10,000 xg for 30 min. Cleared lysates were incubated with Ni-NTA resin (Qiagen) at 4°C on a roller shaker. Lysates were loaded on Poly-Prep chromatography columns (BioRad) and flow-through fractions were collected. Columns were washed twice in wash buffer (50 mM NaH_2_PO_4_, 300 mM NaCl, 20 mM imidazole at pH = 8.0). Beads were resuspended in elution buffer (50 mM NaH_2_PO_4_, 300 mM NaCl, 250 mM imidazole at pH = 8.0). For all the collected fractions (input, flow-through, beads), samples were taken for western blot analysis and leftover were used for RNA extraction using phenol-chloroform. Protein samples were analyzed by western blotting and RNA samples by northern blotting.

For CsrA-His_10_ purification, 100 ml cultures were harvested 4 h following induction in exponential phase. Pellets were lysed in native lysis buffer, 1 mg/ml lysozyme and 10 mM PMSF for 30 min on ice. After sonication and disruption of genomic DNA as described before, lysates were cleared by centrifugation. CsrA-His_10_ was then purified as described in the section on co-immunoprecipitation and using the Qiaexpressionist handbook. All fractions were analyzed by SDS-PAGE. Fractions containing CsrA-His_10_ were further confirmed by western blotting using mouse anti-His antibody (Protein Tech). Dialysis was performed overnight at 4°C in dialysis buffer (50 mM Tris-HCl pH = 8.0, 150 mM NaCl, 10% glycerol, 1mM DTT) and using Spectrum Spectra/Por 3 RC dialysis membrane tubing. Protein yield was quantified using Pierce BCA protein assay kit (Thermo Fisher Scientific).

##### Western blotting

For protein expression, purification and co-immunoprecipitation experiments, 5 μl of cell suspension or fractions were mixed with equal volume 2x Laemmli buffer and heated at 95°C for 5 min then loaded on 10-15% SDS-PAGE gels. For RFP assays, 30 μg total cleared cell lysates were mixed with 4x Laemmli buffer, boiled and loaded on SDS-gels. Protein samples were transferred to a nitrocellulose membrane (Sigma) using a semi-dry blotter (V20-SDB, Scie-Plas). Membranes were blocked in 1 x PBSTM (1x PBS, 0.1% tween, 5% nonfat milk) for 1 h at room temperature on a shaker. Membranes were then incubated with the primary antibody prepared in 1 x PBST in 1:5,000 dilution, overnight at 4°C on a rotary shaker. Membranes were washed three times in 1xPBST then incubated for 1h at room temperature with Licor anti-mouse antibody (1:20,000). After three washes, membranes were dried then scanned with a Licor scanner and visualized using ImageQuant software.

##### Crosslinking experiment

*In vitro* transcribed *fimR2* sRNA containing 4-thio-Us was denatured at 95°C for 2 min then allowed to refold at room temperature for 10 min. The sRNA was then incubated with 30 μg total cell lysates extracted and quantified as described for protein extraction. Incubation was done at 37°C for 30 min. Samples destined for crosslinking were incubated on ice for 15 min under a UV lamp (365 nm). Control samples were incubated on ice without exposure to light. Proteinase K (Roth) treatment was done at 37°C for 15 min. All samples were heated at 95°C for 5 min then loaded on an SDS-PAGE for analysis. The gel was stained in a Coomassie solution, destained in destaining buffer (50% methanol, 10% acetic acid), and then dried on Whatman paper and exposed to a phosphor imaging screen overnight.

##### RNA EMSA

EMSA experiments were performed as described in Yakhnin et al. (2012). *fimR2* 5’-end labelled RNA was heated at 85°C for 3 min then allowed to refold for 10 min at room temperature. RNA was then incubated with or without CsrA-His_10_ in binding buffer (100 mM Tris-HCl pH = 7.5, 100 mM MgCl_2_, 1M KCl), and reaction mixture (7.5% glycerol, 0.2 μg yeast tRNA, 0.5% bromophenol native dye, 20 mM DTT, 0.04 U RNasin Plus RNase inhibitor). lCsrA-His_10_ was diluted in dilution buffer (10 mM Tris-HCl pH = 7.5, 2 mM DTT, 10% glycerol). Reactions were incubated at 37°C for 30 min then loaded on a 10% native gel (10% acrylamide, 2.5% glycerol, 0.5x TBE) and run for 1 h at 200 V. The gel was then wrapped in plastic foil and exposed to a phosphor imaging screen at -20°C.

##### RACE

5’- and 3’-RACE experiments were conducted using the FirstChoice™ RLM-RACE Kit (Invitrogen) and as per manual description. Twenty micrograms of total RNA were separated on an 8% denaturing polyacrylamide gel and using Riboruler LR RNA ladder (Thermo Fisher Scientific), gel pieces corresponding to RNA bands in the range of 20-50 nucleotides were cut, crushed, and eluted overnight at 4°C in elution buffer (0.3 M sodium acetate pH = 5.5, 1 mM EDTA). On the following day, the supernatant was precipitated in 100% ethanol. RNA was then poly-adenylated using *E. coli* Poly(A)-Polymerase (New England Biolabs) as suggested at 37°C for 30 min. RNA was then purified from reaction components by a phenol-chloroform extraction. For 5’-RACE experiments, 5’-RACE adapter was annealed to template RNA using RNA Ligase (Thermo Fisher Scientific). cDNA synthesis was carried out using SuperScript IV reverse transcriptase (Thermo Fisher Scientific) with random hexamer primer (Thermo Fisher Scientific) with 10-min cycles at 23°C, 55°C, and 80°C. For 3’-RACE experiments, 3’-RACE adapter was used as a primer for cDNA synthesis using Superscript IV with 10-min cycles at 55°C and 80°C. *fimR2S*-specific primers were then used in combination with 5’-RACE and 3’-RACE outer primers in a first amplification step, and 5’-RACE and 3’-RACE inner primers in a second amplification step. PCR reactions were carried out using a homemade Taq Polymerase. PCR products were purified using Wizard^®^ SV Gel and PCR Clean-Up (Promega) and ligated into the pGEM-T-Easy vector (Promega) using supplied DNA Ligase. Ligated products were transformed into XL1-Blue competent cells. Plasmid extraction was done with Wizard^®^ Plus SV Minipreps DNA Purification System (Promega). Inserts were sequenced using T7 universal primer from Microsynth.

##### RT-qPCR

RT-qPCR was conducted as described (Raad et al., 2021). One microgram of total RNA from all experimental conditions were treated with DNase I (Thermo Fisher Scientific) to digest DNA, according to the manufacturer’s protocol. Samples were then reverse transcribed into cDNA with SuperScript™ IV One-Step RT-PCR System (Invitrogen) and random primer hexamers (Thermo Fischer Scientific). After reverse transcription, cDNA samples were treated with RNase H (NEB) to hydrolyze leftover RNA. qPCR was done using GoTaq^®^ qPCR Master Mix (Promega), 50-fold diluted cDNA, and a final concentration of 250 nM to 1 M of oligonucleotides. All primer pairs used for qPCR analysis were optimized using a standard curve. qPCR reactions were prepared by the CAS-1200 Corbett robot (Corbett Robotics) and analyzed using the Rotor Gene 6000, with suggested standard cycling conditions. *recA* was used as an internal control for the normalization of gene expression. All three biological replicates used for this analysis were run in duplicates. The 2^−ΔΔCT^ method was used to calculate the fold-change relative to the control (Rao et al., 2013). The mean log_2_ fold-change and standard error of the mean were computed.

##### PCR

To confirm that the deletion strains acquired from the KEIO collection contain a kanamycin cassette instead of the ORF of the deleted locus, PCR amplification was done using oligonucleotides K1 and K2 combined with gene-specific primers (See **Table S2**) as suggested in Datsenko and Wanner (2000), and using home-made Taq Polymerase. For genomic DNA preparation, 1.5 mL of bacterial culture were lysed with 1.2 mg/mL Proteinase K (Roth) and 6% SDS in TE buffer (10 mM Tris-HCl pH = 8.0 and 1 mM EDTA), at 37°C for 1 h. Genomic DNA was then extracted with basic phenol-chloroform extraction and precipitated in 2.5 volumes of 100% ethanol and 0.3 M sodium acetate.

##### Fluorescence assay

For RFP experiments, 500 μl of cultures, in 2 technical replicates, and 3 biological replicates were pipetted into 48-well plate (cellstar) following induction with IPTG and Tetracyclin. Fluorescence and OD_600_ were measured using Tecan plate reader with RFP excitation at 584 nm and emission at 607 nm. Background fluorescence from bacterial strains with sRNA overexpression plasmids and target-RFP expression plasmids were subtracted from experimental data.

For sicA-GFP, analysis was done similarly and GFP fluorescence was measured using Tecan plate reader with GFP excitation at 488 nm and emission at 525 nm. As *fimR2S-S* overexpression does not cause biofilm formation, fluorescence measurements were divided over OD_600_ of cultures (also measured using Tecan plate reader) to obtain absolute fluorescence. Background fluorescence from SB300 cells normalized over OD_600_ was subtracted from experimental samples.

##### Biofilm assay

Biofilms assays were conducted as described in Merritt et al. (2005) with minor modifications. Bacterial strains were diluted 1:100 from overnight cultures into fresh LB medium. OD_600_ = 0.4, 200 μl of culture were inoculated in 96-well flat bottom microtiter plates (cellstar) for 2 h at 37°C (for exponential phase samples) and 22 h (for stationary phase samples). Planktonic bacteria were then removed by two washes with H_2_O. Biofilms were then stained with 0.1% crystal violet in water for 15 min at room temperature. Wells were washed three times with H_2_O and crystal violet was solubilized with a solution of 80% Ethanol and 20% Acetone. OD_600_ of solubilized crystal violet was measured using Vmax microplate reader (LabX Molecular Devices).

For micrographs, 500 μl of cultures were inoculated in 12-well flat bottom plates (TPP) mounted with a coverslip and incubated as indicated for liquid culture-based biofilm assays. Coverslips were then washed with H_2_O and stained with 0.1% crystal violet as described. Dried coverslips were mounted on glass microscope slides and visualized using a Leica Microscope.

##### Scanning Electron Microscopy

For visualizing bacterial cells in suspension by scanning electron microscopy, 150 μl of bacterial cells were taken from growing cultures 2 h after reaching OD_600_ = 0.4 (exponential phase) and 22 h post-inoculation (stationary phase). For inspecting biofilm potential, 1 mL of bacterial cells were inoculated in 24-well plate (TPP) mounted with 12 mm coverslips at OD_600_ = 0.4. Plates were incubated at 37°C for 2 h (exponential phase) or 22 h (stationary phase), under static conditions. Bacterial cell fixation was done as described in (Winzer et al., 2020). Briefly, cell pellets and coverslips were washed with 0.1 M sodium cacodylate (pH = 7.2) and fixed in 2% glutaraldehyde in sodium cacodylate buffer for 2 h at room temperature. Washes were done with 0.1 M sodium cacodylate and samples were further fixed with 2% osmium tetroxide in sodium cacodylate for 2 h at room temperature. After washing with water, the samples were dehydrated in increasing concentrations of ethanol (30%-100%). Samples were then resuspended or immersed in hexamethyl-disilazane and dried on metal SEM holders. Samples were sputter-coated with gold and visualized with a Zeiss Gemini 450 microscope operating at 5kV.

##### HeLa infection assay

HeLa invasion assays were carried out as described in (Pfister et al., 2020). Bacterial inocula for infection were cultured in LB medium with 0.3 M NaCl from overnight cultures at 37°C until OD_550_ = 0.6. Induction of SB300 strains containing pBbE6k-*fimRS2-S* was done with IPTG at OD_550_ = 0.4. Stationary phase samples were taken 24 h post-inoculation. HeLa (Kyoto) cells were seeded into 24-well plates (TPP) and grown in Dulbecco modified Eagle’s medium (DMEM) supplemented with 10% fetal bovine serum (FBS) for 24 h at 37°C. Before infection with SB300 cells and derivatives, adherent cells were washed then incubated in Hank’s buffered salt (HBSS) for 10 min. Infections were carried out using 3 x 106 CFU of bacteria per well, for 50 min at 37°C. Extracellular bacteria were inactivated by the addition of DMEM containing 10% FBS and gentamicin (400 μg/mL) and subsequent incubation at 37°C for 30 min. HeLa cells were then washed in PBS and lysed in 0.1% sodium-deoxycholate in PBS to release intracellular bacteria. The initial bacterial inocula and intracellular loads were quantified on LB/agar plates containing Streptomycin and/or Kanamycin.

##### CopraRNA Analysis

The analysis was done using default parameters of CopraRNA and with a search for target interactions 300 nucleotides upstream and downstream mRNA start codon. As an input, the first 18 nucleotides of the *fimR2* sequence were used (5’-UUCAGGGACGUCAUUACG-3’).

##### MFold prediction

*fimR2* secondary structure was predicted using MFold algorithm (Zuker, 2003) with default values used for all parameters. The structure with the lowest predicted free energy is shown in this study.

##### Statistical analysis

Unpaired two-tailed t-test was used with Welsh’s correction for all significance analysis and was conducted in Graphpad Prism. All analysis p-values are indicated in the figure legends.

## Supplemental Information title and legends

**Table S1: List of strains and plasmids.**

**Table S2: List of oligonucleotides.**

