## Supplemental Figures for "The stationary phase-specific sRNA *fimR2* is a multifunctional regulator of bacterial motility, biofilm formation and virulence"

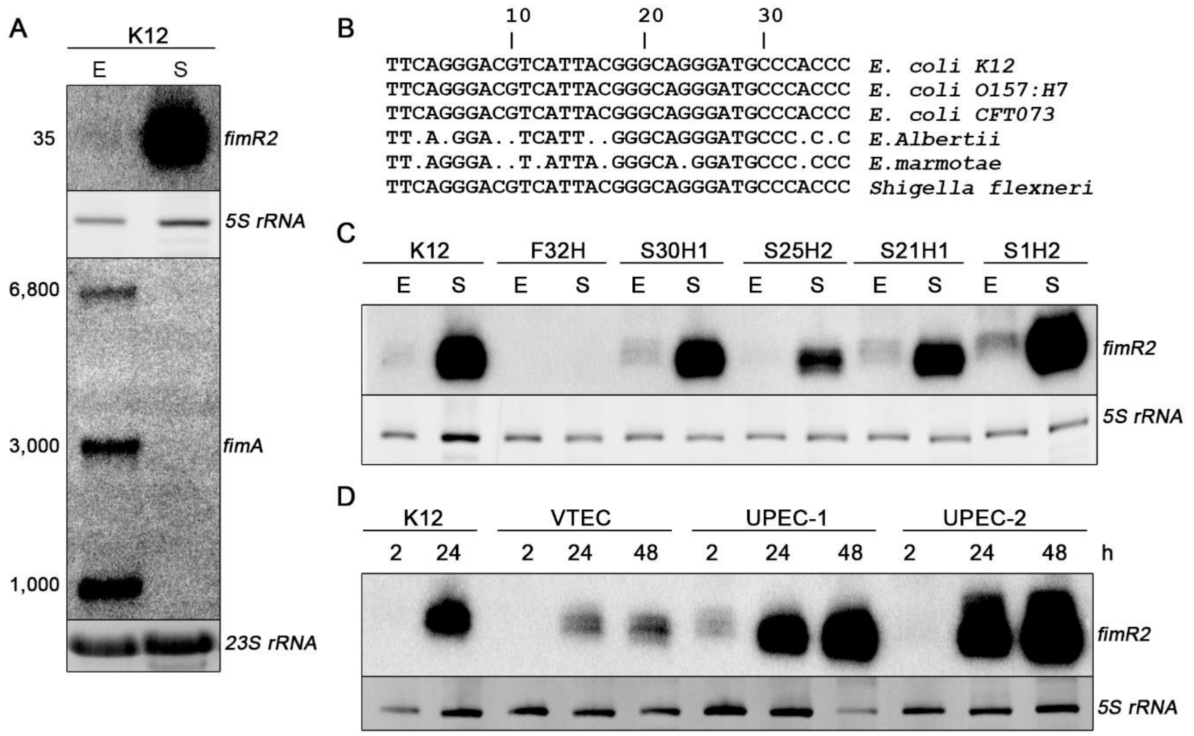


**Figure S1: *fimR2* is expressed in various *E. coli* strains**. **A.** Northern blot showing *fimR2* and *fimAICDFGH* expression in the K12 *E. coli* strain in exponential (E) and stationary (S) phase, respectively. Ethidium bromide staining of *5S rRNA* and *23S rRNA* are shown as loading controls on an 8% denaturing polyacrylamide gel and 1.2% agarose gel, respectively. Estimated transcript sizes are shown on the left in nucleotides. **B.** Alignment of the *fimR2* sequences from various enterobacterial strains. Sequential numbers indicate nucleotide positions. Dots indicate nucleotide mismatches compared to the K12 strain. **C.** Northern blot showing *fimR2*-phase dependent expression in ESBL *E. coli* strains. Total RNA samples from E (exponential phase) and S (stationary phase) are shown. Ethidium bromide staining of *5S rRNA* is shown as a loading control**. D.** Northern blot analysis of *fimR2* expression in VTEC, UPEC-1, and UPEC-2 strains. Total RNA samples from different time points of bacterial growth, 2, 24, and 48 h (hours), are shown. Ethidium bromide staining of *5S rRNA* is shown as a loading control**.**

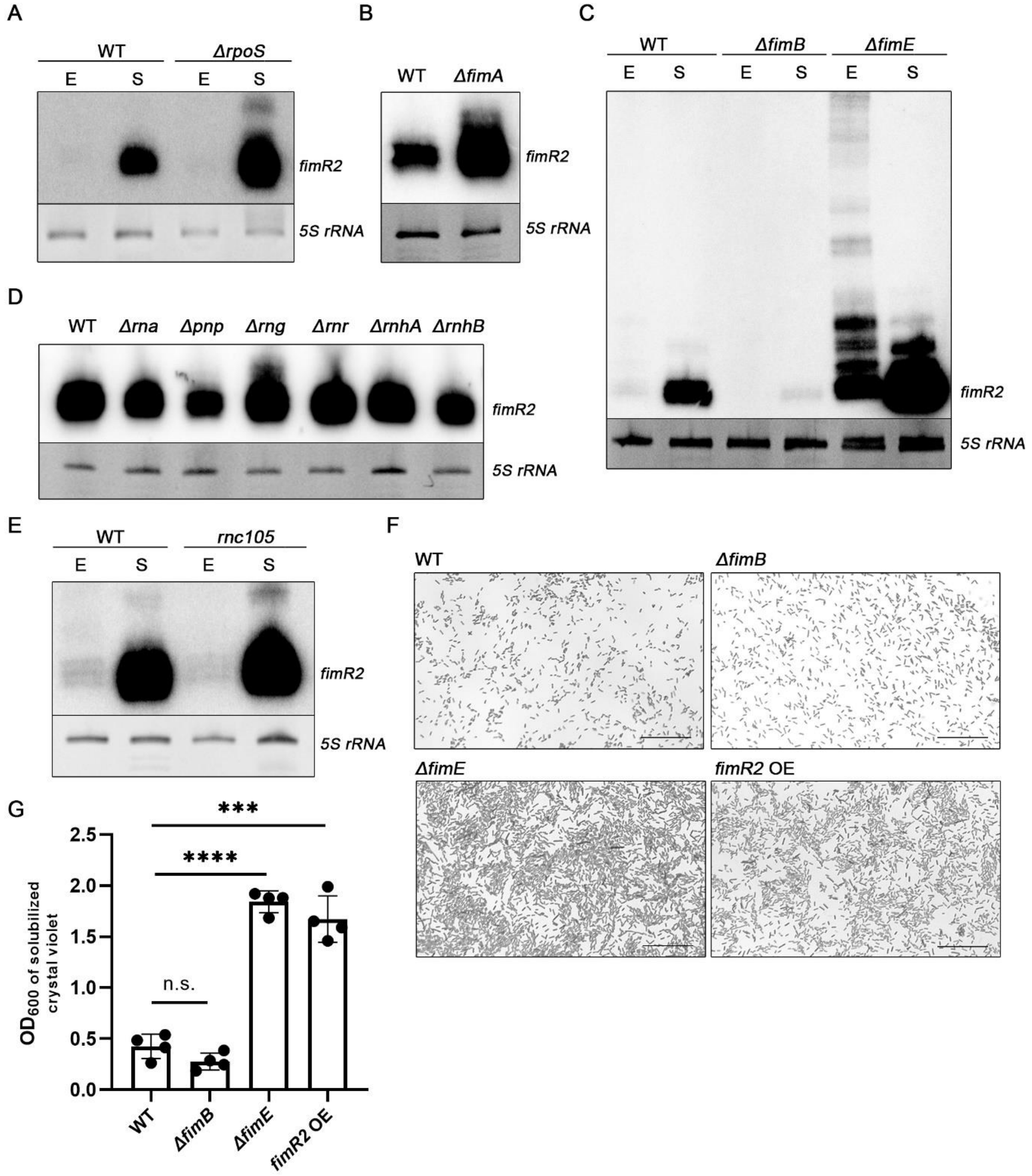


**Figure S2:** ***fimR2*** **expression is** ***fimAICDFGH*****-dependent. A.** Northern blot showing *fimR2* expression in WT–and *Δrpos* (*rpoS* deletion). RNA samples from E (exponential phase) and S (stationary phase) are shown. Ethidium bromide staining of *5S rRNA* is shown as a loading control. **B.** Northern blot analysis of *fimR2* expression in WT and *ΔfimA* (*fimA* deletion). Total RNA samples from stationary phase are shown. Ethidium bromide staining of *5S rRNA* is shown as a loading control. **C.** Northern blot analysis of *fimR2* expression in WT, *ΔfimB* (*fimB* deletion), and *ΔfimE* (*fimE* deletion) strains**.** RNA samples from E (exponential phase) and S (stationary phase) are shown. Ethidium bromide staining of *5S rRNA* is shown as a loading control. **D.** Northern blot showing *fimR2* expression in WT, *Δrna* (RNase I deletion), *Δpnp* (PNPase deletion), *Δrng* (RNase G deletion), *Δrnr* (RNase R deletion), *ΔrnhA* (RNase HI deletion), and *ΔrnhB* (RNase HII deletion) strains. Total RNA samples from stationary phase are shown. Ethidium bromide staining of *5S rRNA* is shown as a loading control**. E.** Northern blot showing *fimR2* expression in WT and *rnc105* (RNase III mutant) strains. RNA samples from E (exponential phase) and S (stationary phase) are shown. Ethidium bromide staining of *5S rRNA* is shown as a loading control. **F.** Micrographs of air-liquid phase biofilms stained with crystal violet, under conditions from C**.** Scale bar = 25 μm. **G.** Quantitative biofilm assay showing mean + SD $OD_{600}$ of crystal violet staining from four biological replicates of strains in C**.** Unpaired two-tailed t-test with Welch's correction was used to determine significance with n.s and * showing not significant and significant results, respectively. The p-values are in order 0.0916, <0.0001, and 0.0003.

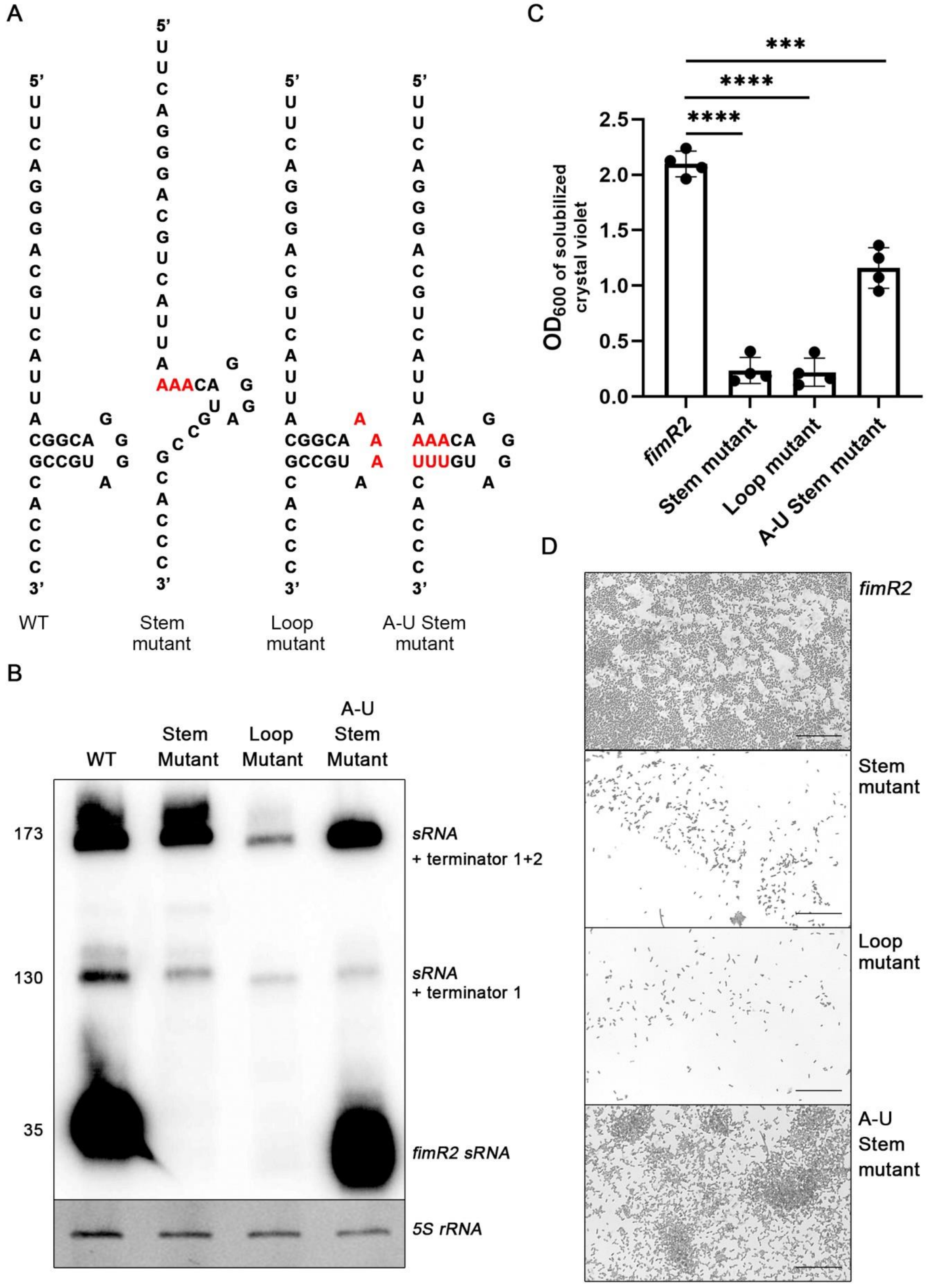


**Figure S3: *fimR2* stem-loop is important for sRNA stability**. **A.** MFold predicted secondary structure of *fimR2* and mutant constructs. Point mutations are indicated in red. **B.** Northern blot of *fimR2* and *fimR2* mutants following overexpression of sRNAs in exponential phase. Ethidium bromide staining of *5S rRNA* is shown as a loading control. Sizes are indicated on the left in nucleotides. **C.** Quantitative biofilm assay showing mean + SD of $OD_{600}$ of crystal violet-stained biofilms from four biological replicates. Unpaired two-tailed t-test with Welch's correction was used to determine significance (*). The p-values are in order <0.0001, <0.0001, and 0.0003. **E.** Micrographs of air-liquid phase biofilms stained with crystal violet, following overexpression of *fimR2* and *fimR2* mutants. Scale bar = 25 μm.

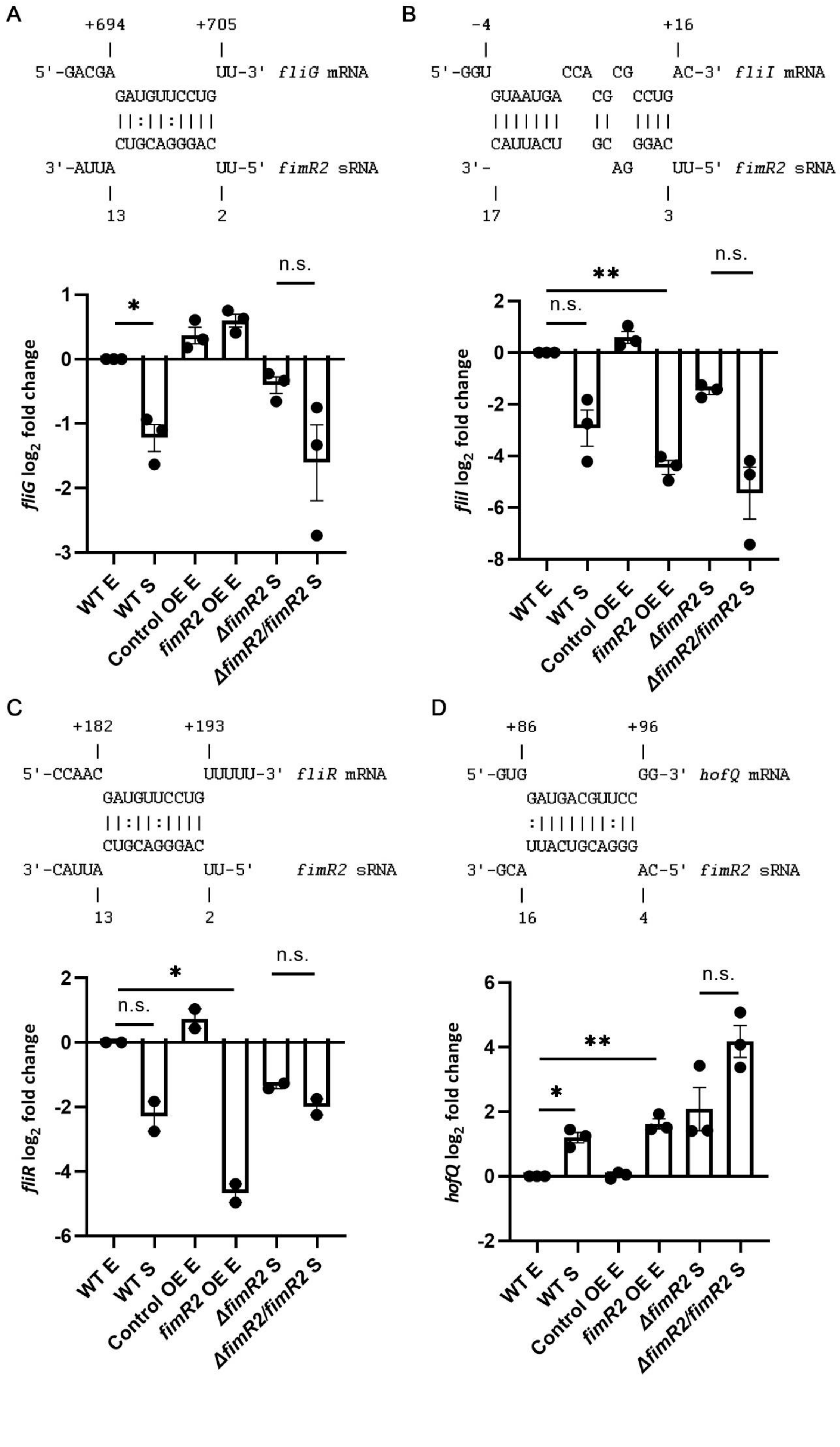
A
+694
+705
5'-GACGA
UU-3' fliG mRNA
GAUGUUCCUG
||:||:||||
CUGCAGGGAC
3'-AUUA
UU-5' fimR2 sRNA
13
2
B
-4
+16
5'-GGU
CCA CG
AC-3' fliI mRNA
GUAAUGA CG CCUG
||||||| || ||||
CAUUACU GC GGAC
3'-
AG
UU-5' fimR2 sRNA
17
3
fliG log2 fold change
fliI log2 fold change
n.s.
*
**
WT E
WT S
Control OE E
fimR2 OE E
ΔfimR2 S
ΔfimR2/fimR2 S
C
+182
+193
5'-CCAAC
UUUUU-3' fliR mRNA
GAUGUUCCUG
||:||:||||
CUGCAGGGAC
3'-CAUUA
UU-5' fimR2 sRNA
13
2
D
+86
+96
5'-GUG
GG-3' hofQ mRNA
GAUGACGUUCC
:|||||||:||
UUACUGCAGGG
3'-GCA
AC-5' fimR2 sRNA
16
4
fliR log2 fold change
hofQ log2 fold change

**Figure S4: *fimR2* affects the expression of several target mRNAs.** Predicted interaction site with *fimR2* (top) and RT-qPCR analysis (bottom) of **A.** *fliG,* **B.** *fliI,* **C.** *fliR* and **D.** *hofQ* mRNAs in WT, Control OE (control overexpression), *fimR2* OE (*fimR2* overexpression), *ΔfimR2* (*fimR2* deletion), and *ΔfimR2*/*fimR2* (*fimR2* complementation) strains. Samples from E (exponential phase) and S (stationary phase) are shown. Mean $\log_2$ fold change + SEM are shown for all candidates from three biological replicates. $\log_2$ fold change was based on comparison with WT E samples. Unpaired two-tailed t-test with Welch's correction was used to determine significance with n.s and * showing not significant and significant results, respectively. The p-values resulting from this analysis are computed in comparison with WT E and *ΔfimR2* samples and are in order **A.** 0.0282 and 0.1733, **B.** 0.0527, 0.0037, and 0.05757, **C.** 0.1270, 0.0395, and 0.2074, and **D.** 0.0175, 0.0084, and 0.0713.

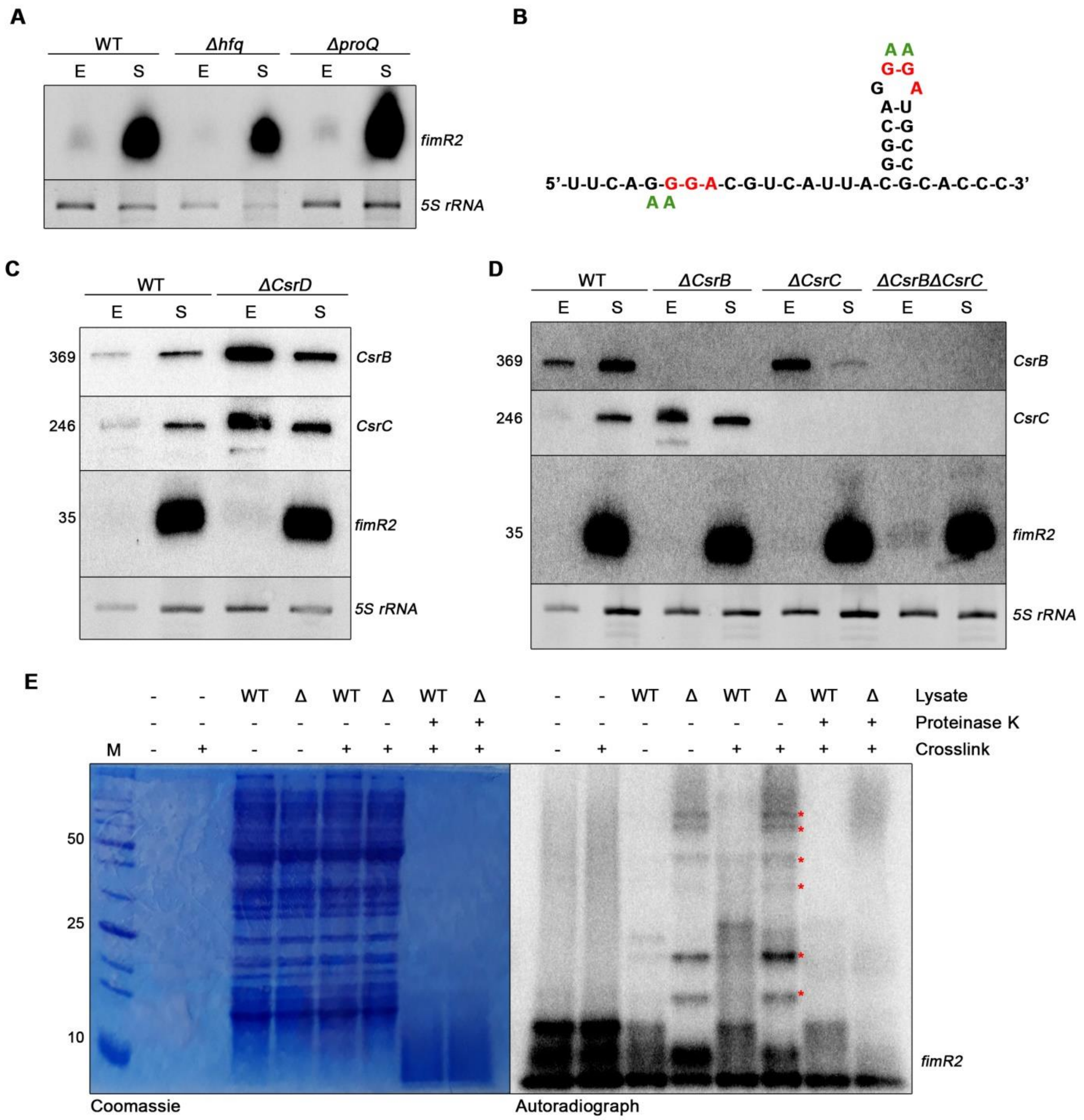


**Figure S5: *fimR2* is an Hfq/ProQ-independent sRNA. A.** Northern blot of *fimR2* expression in WT, *Δhfq* (*hfq* deletion), *ΔproQ* (*proQ* deletion) strains. Total RNA samples from E (exponential phase) and S (stationary phase) are shown. Ethidium bromide staining of *5S rRNA* is shown as a loading control. **B.** MFold-predicted secondary structure of *fimR2* with putative CsrA binding sites indicated in red. Nucleotides in green represent substitution mutations. **C.** Northern blot analysis of *fimR2*, *CsrB*, and *CsrC* in WT and *ΔCsrD* (*CsrD* deletion). Total RNA samples from E (exponential phase) and S (stationary phase) are shown. Ethidium bromide staining of *5S rRNA* is shown as a loading control. sRNA sizes are indicated on the left in nucleotides. **D.** Northern blot analysis of *fimR2*, *CsrB*, and *CsrC* in WT, *ΔCsrB* (*CsrB*

deletion), *ΔCsrC* (*CsrC* deletion), and *ΔCsrBΔCsrC* (*CsrB* and *CsrC* double deletion). Total RNA samples from E (exponential phase) and S (stationary phase) are shown. Ethidium bromide staining of *5S rRNA* is shown as a loading control**.** sRNA sizes are indicated on the left in nucleotides. **E.** Coomassie staining (left) of total cell lysates and autoradiograph (right) of 5'-end labelled *fimR2 in vitro* transcribed with 4-thiouridine incubated with total cell lysates from WT and Δ (*fimR2* deletion), + (with) and – (without) crosslinking by ultraviolet light (wavelength of 365nm) and treatment with Proteinase K. M indicates the lane with the unstained protein marker. The sizes of prominent marker bands are indicated on the left in kDa. Upshifts are marked with red asterisks.

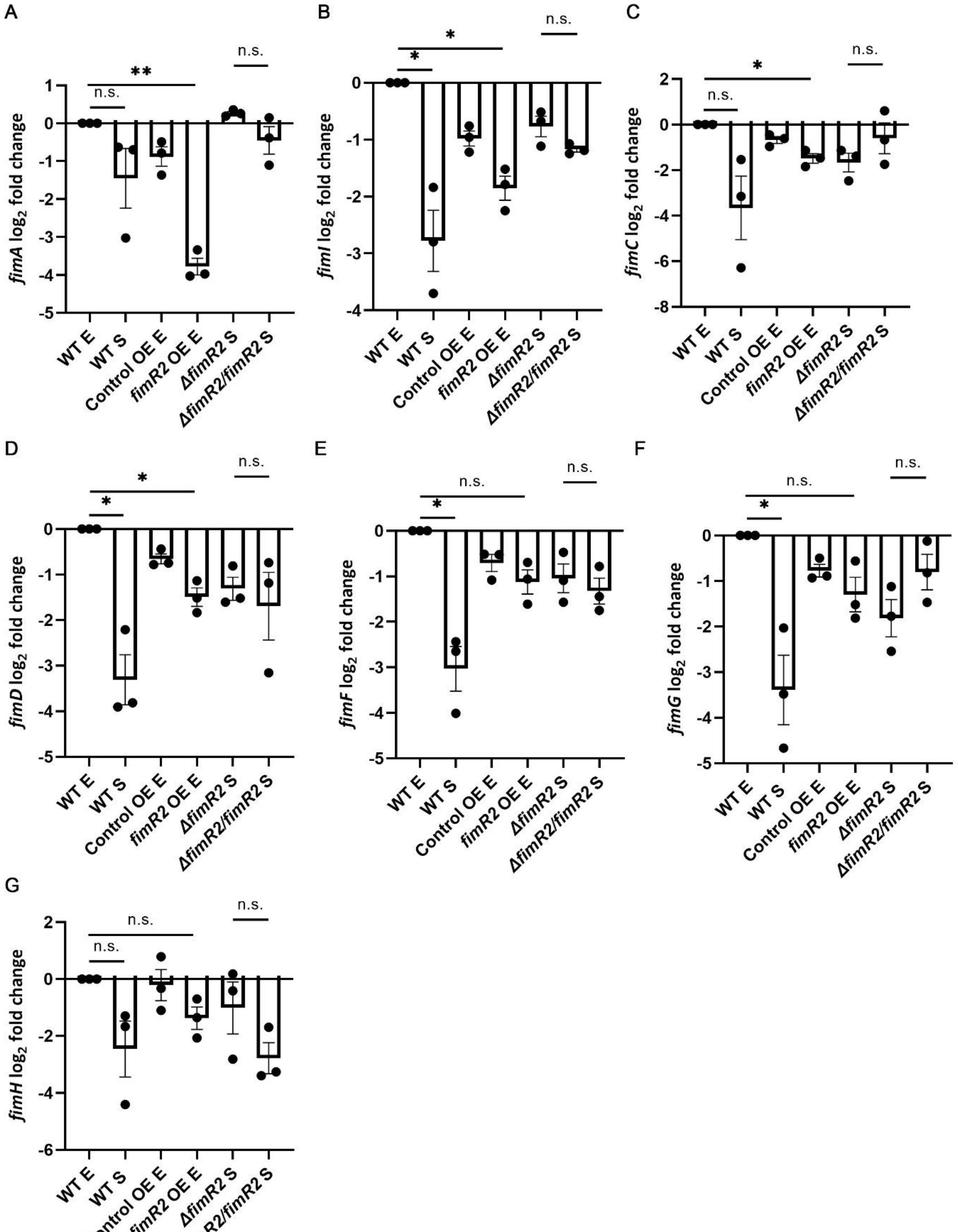


**Figure S6: *fimR2* downregulates *fimAICDFGH*.** RT-qPCR analysis of **A.** *fimA,* **B.** *fimI,* **C.** *fimC,* **D.** *fimD,* **E.** *fimF,* **F.** *fimG,* and **G.** *fimH* in WT, Control OE, *fimR2* OE, *ΔfimR2*, and *ΔfimR2/fimR2* strains. Total RNA samples from E (exponential phase) and S (stationary phase) are shown. Mean $\log_2$ fold change + SEM are shown for all seven transcripts from three biological replicates. $log_2$ fold change was based on comparison with WT E samples. Unpaired two-tailed t-test with Welch's correction was used to determine significance with n.s and * showing not significant and significant results, respectively. The p-values in comparison with WT E or *ΔfimR2* samples are **A.** 0.2064, 0.0034, and 0.1852, **B.** 0.0354, 0.0130, and 0.1471, **C.** 0.1198, 0.0185, and 0.2652**, D.** 0.0266, 0.0178, and 0.6663, **E.** 0.0254, 0.0519, and 0.5482, **F.** 0.0471, 0.0756, and 0.1486, **G.** 0.1293, 0.0731, and 0.1883.
